# scDiagnostics: systematic assessment of cell type annotation in single-cell transcriptomics data

**DOI:** 10.64898/2026.01.29.701618

**Authors:** Anthony Christidis, Andrew Ghazi, Smriti Chawla, Nitesh Turaga, Robert Gentleman, Ludwig Geistlinger

## Abstract

Although cell type annotation has become an integral part of single-cell analysis workflows, the assessment of computational annotations remains challenging. Many annotation tools transfer labels from an annotated reference dataset to a new query dataset of interest, but blindly transferring labels from one dataset to another has its own set of challenges. Often enough there is no perfect alignment between datasets, especially when transferring annotations from a healthy reference atlas for the discovery of disease states. We present `scDiagnostics`, a new open-source software package that facilitates the detection of complex or ambiguous annotation cases that may otherwise go unnoticed, thus addressing a critical unmet need in current single-cell analysis workflows. `scDiagnostics` is equipped with novel diagnostic methods that are compatible with all major cell type annotation tools. We demonstrate that `scDiagnostics` reliably detects complex or conflicting annotations using both carefully designed simulated datasets and diverse real-world single-cell datasets. Our evaluation demonstrates that `scDiagnostics` reliably identifies misleading annotations that systematically distort downstream analysis and interpretation and that would otherwise remain undetected. The `scDiagnostics` R package is available from Bioconductor (https://bioconductor.org/packages/scDiagnostics).

## Background

Single-cell RNA sequencing (scRNA-seq) has transformed our ability to profile transcriptomic diversity at the level of individual cells, enabling unprecedented insights into cellular heterogeneity within complex tissues and biological systems [1, 2]. A fundamental and critical step in any scRNA-seq analysis workflow is the accurate identification and annotation of cell types, which provides essential biological context for understanding cellular functions, developmental trajectories, and disease mechanisms [3, 4]. This process serves as the foundation for virtually all downstream analyses, from cell-cell interaction studies to differential expression analysis and pathway enrichment, making the quality of cell type annotations paramount to the biological interpretability and validity of research findings [5, 6].

Traditional approaches to cell type annotation have relied heavily on manual curation by domain experts, who examine gene expression patterns within computationally derived cell clusters and compare them against known cell type-specific marker genes [1, 2]. Although this manual process can achieve high accuracy when performed by experienced researchers, it is inherently labor intensive, time-consuming, subjective, and does not scale well to the increasingly large datasets generated by modern single-cell technologies. Moreover, manual annotation suffers from reproducibility challenges, as different experts may reach different conclusions when interpreting the same data.

To address these limitations, the field has witnessed the rapid development of computational approaches for automated cell type annotation [7]. These methods can be broadly categorized into two theoretical frameworks: (1) marker gene-based approaches that leverage curated databases of cell type-specific genes to score and classify cells based on the expression of known biomarkers [8, 9], and (2) reference-based methods that compare single-cell expression profiles against well-annotated reference datasets, transferring labels based on transcriptomic similarity [10, 11]. Numerous computational tools implementing both of these methodological strategies have been developed and have contributed to significant advances in the field, including `SingleR` [12] for reference-based annotation using correlation analysis, `CellTypist` [13] for supervised classification trained on labeled reference data, CHETAH [14] for hierarchical classification with confidence scoring, `scmap` [15] for both cluster-based and cell-based annotation, `Seurat’s` [16] integrated label transfer framework `Azimuth` [17], and more recent deep learning approaches like `scDeepSort` [18] using graph neural networks and `scArches` [19] for transfer learning across datasets.

Despite these advances, several fundamental challenges persist in the current landscape of computational cell type annotation [6]. Most critically, many of these tools operate as “black boxes” where the decision-making process is opaque to users. Researchers often apply these methods without sufficient understanding of their underlying assumptions, limitations, or the quality of the resulting annotations. This black-box nature is particularly problematic because annotation errors can propagate through entire analysis pipelines, potentially leading to incorrect biological conclusions.

A central challenge in reference-based annotation is the label transfer problem, where annotations from a reference dataset are computationally projected onto a new query dataset of interest [10, 12, 13, 19]. Although conceptually straightforward, this process is fraught with difficulties. Reference and query datasets often exhibit technical batch effects arising from differences in sequencing platforms, experimental protocols, library preparation methods, or laboratory conditions. Additionally, biological differences between datasets—such as differences in tissue sampling, cell culture conditions, developmental stages, or disease states—can create misalignments that compromise annotation accuracy.

Perhaps most critically, there is frequently no perfect alignment between reference and query datasets, particularly when transferring annotations from healthy reference atlases to study disease states or novel biological conditions [20]. In such scenarios, cells in the query dataset may represent cellular states that are not well-represented in the reference, leading to forced assignments to the most similar available cell type rather than appropriate identification of novel or intermediate cell states. This problem is exacerbated when dealing with rare cell types [21], transitional cell states [22], or cells undergoing dynamic biological processes [23].

This gap in quality assessment is particularly problematic because complex or ambiguous annotation cases often go unnoticed in standard workflows. Cells that fall on a developmental continuum, exhibit intermediate expression profiles between known cell types, or represent novel cellular states may be incorrectly forced into existing categories without appropriate flagging for manual review. Similarly, systematic biases in annotation—such as the tendency of methods to favor well-represented cell types over rare populations—may not be detected without proper diagnostic tools.

Although several tools exist that can be used for the diagnosis of such annotation failures, these approaches are typically task-specific and not generally applicable. This includes cell type annotation tools providing classification confidence scores [12, 17], deep-learning models for out-of-distribution (OOD) detection [19], and specific tools for identifying disease-driven perturbations [20]. For instance, deep-learning OOD detection tools typically require users to adopt their specific neural network architectures, precluding their use for diagnostic audits in established pipelines. Consequently, users typically lack tool-agnostic systematic diagnostics to explore and evaluate whether their reference dataset is appropriate for their query, whether the annotation transfer was successful, and whether specific cell populations may have been misannotated.

We present `scDiagnostics`, a new open-source software package that facilitates the detection of complex or ambiguous annotation cases that may otherwise go unnoticed, thus addressing a critical unmet need in current single-cell analysis workflows. Our package introduces new standards of rigor to cell type annotations derived from automated reference mapping, ensuring more accurate and reliable cell type annotations, leading to clearer and more dependable biological insights.

## Results

### Overview of the scDiagnostics package

The `scDiagnostics` R/Bioconductor package implements functionality to perform a systematic quality assessment of cell type annotations through statistical comparison of query and reference datasets (Fig. 1). The package is agnostic to the annotation method used and is applicable to annotations derived from manual marker gene-based approaches (Fig. 1A, top) and automated reference-based label transfer (Fig. 1A, bottom). The package requires as inputs: (i) a query dataset with cell type predictions, and (ii) an annotated reference dataset representing the biological system or state of interest. However, even with expert-annotated reference atlases, dataset incompatibilities can lead to misclassifications and undetected novel cell states (Fig. 1B). Although popular cell type annotation tools provide useful per-cell confidence or prediction scores that quantify annotation uncertainty, relying on these metrics alone has several limitations and does not systematically diagnose the underlying causes of low confidence assignments (Supplementary Methods S1.1–S1.2 and Supplementary Table S1).. The `scDiagnostics` package therefore provides users with functionality to evaluate the compatibility of query and reference data through three complementary analytical components, each designed to verify annotations at the level of the assigned cell type categories (Fig. 1C-E).

**Fig. 1.**
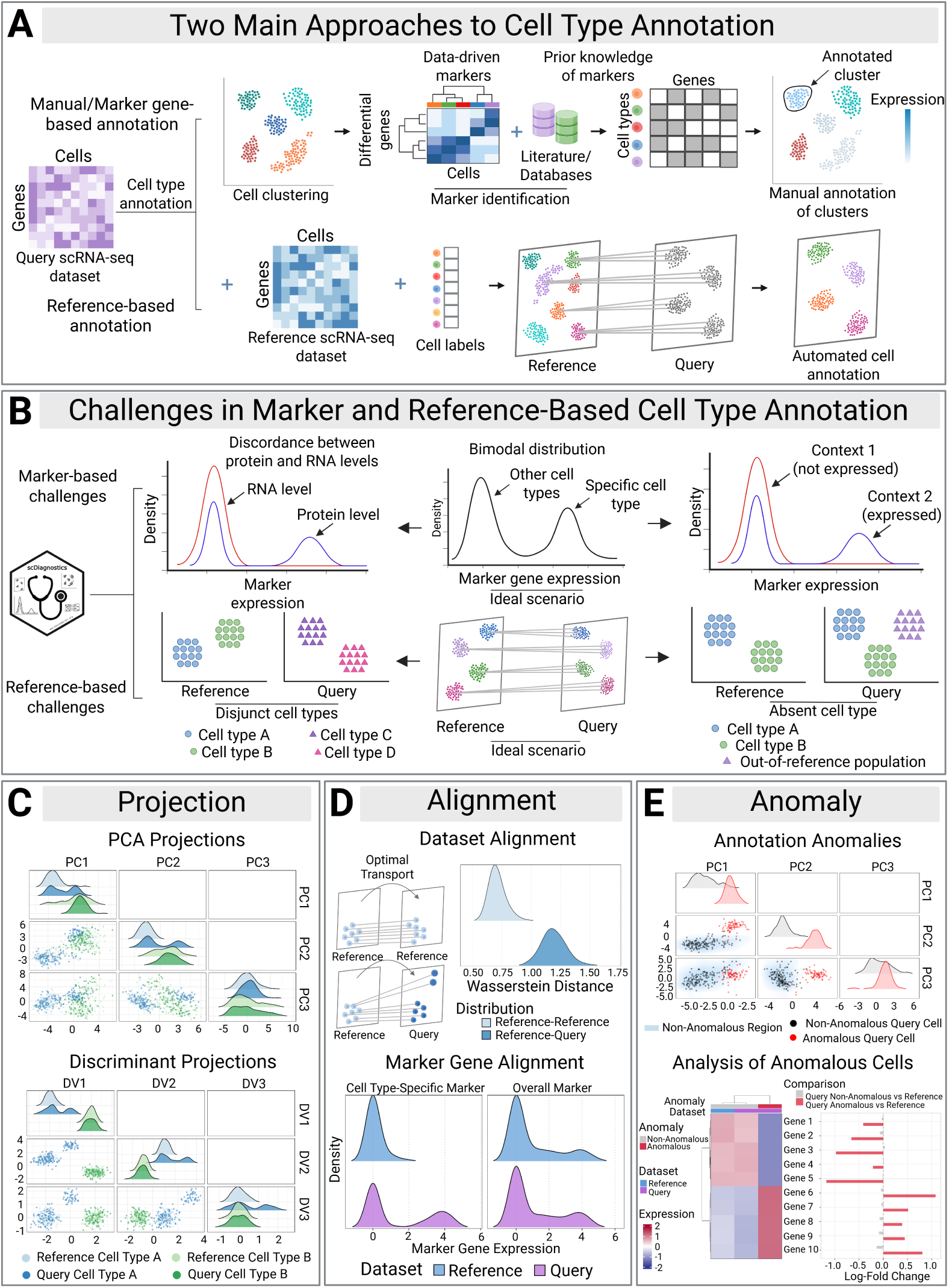
Overview. (**A**) The two main approaches for cell type annotation of single cell data: manual annotation of empirically derived cell clusters using known marker genes (top), and automated annotation transfer from a reference dataset (bottom). (**B**) The challenges associated with manual marker gene-based annotation (top) and reference-based annotation transfer (bottom), including complex gene expression patterns and cell populations that are not shared between reference and query dataset. (**C**) Projection of the query dataset onto a low-dimensional (e.g. PCA or discriminant) space spanned by the reference dataset. (**D**) Evaluation of dataset and marker gene alignment between reference and query datasets using optimal transport and concordance analysis. (**E**) Detection and characterization of annotation anomalies in the query dataset, highlighting the transcriptomically divergent anomalous populations (red).

First, projection functions visualize query cells alongside their assigned reference counterparts within a low-dimensional latent space. In contrast to non-linear visualization techniques that can distort local density [24], `scDiagnostics` uses interpretable, statistically sound projection methods (Fig. 1C). Beyond these projection-based approaches, the package also supports alternative visualization techniques to assess the preservation of pairwise distances between matched reference and query populations (see Methods, *Exploratory data analysis of cell type annotations*).

Second, the alignment component evaluates the similarity between matched reference and query cell types. The package offers a comprehensive suite of functions to determine whether matched cell types display similar expression patterns in a reduced dimensional space based on global distributional metrics (Fig. 1D, top). In addition, the module allows users to assess dataset alignment on the level of individual cell type-specific marker genes (Fig. 1D, bottom).

Third, the anomaly detection component employs outlier detection algorithms to identify specific query cells that deviate significantly from their assigned reference cell type (Fig. 1E, top). By flagging and isolating these anomalous query cells, users can apply established downstream analysis steps—such as differential expression or cell-cell interaction analysis—to further characterize these cells (Fig. 1E, bottom). Rather than simply discarding these cells or conflating them with predefined reference categories, `scDiagnostics` provides analysts with functionality to investigate and distinguish potential causes for the observed anomalies, including (i) *technical artifacts*, such as batch effects, quality-control issues, doublets, or segmentation artifacts; (ii) *reference incompleteness*, representing true out-of-reference populations absent from the reference; (iii) *annotation ambiguity*, reflecting mapping errors or ambiguous annotation despite adequate reference coverage; and (iv) *biological drift*, such as disease activation or trajectory shifts within a known cell type (Supplementary Figure S1).

To facilitate broad adoption and accessibility for applied users, the package is equipped with comprehensive documentation, detailed tutorials, and intuitive visualization methods that present results from the cell type annotation diagnostics in a clear and user-friendly manner (see Methods, *Code availability*). Furthermore, computational benchmarking demonstrates that `scDiagnostics` provides highly performant functionality with minimal runtime and memory requirements. Even for reference atlases with hundreds of thousands of cells, the main diagnostic functions can be executed almost instantaneously, allowing the framework to be applied seamlessly also to large single-cell datasets (Supplementary Results S2.4 and Supplementary Figure S2).

In the following, we evaluate the diagnostic capabilities of `scDiagnostics` using simulated single-cell data with known ground truth composition, and demonstrate its utility in two case studies with real single-cell datasets. These applications show that the package can identify misleading annotations resulting from dataset incompatibilities, facilitate the detection of disease-associated cell states that would otherwise be obscured, and therefore significantly improve downstream analysis and biological interpretation.

### scDiagnostics detects out-of-reference populations in simulated single-cell data

To demonstrate the utility of `scDiagnostics` for detecting misleading annotations in a controlled setting, we generated synthetic scRNA-seq datasets with known ground truth composition. We simulated two different scenarios to mimic common challenges in reference-based cell type annotation (Fig. 2).

**Fig. 2.**
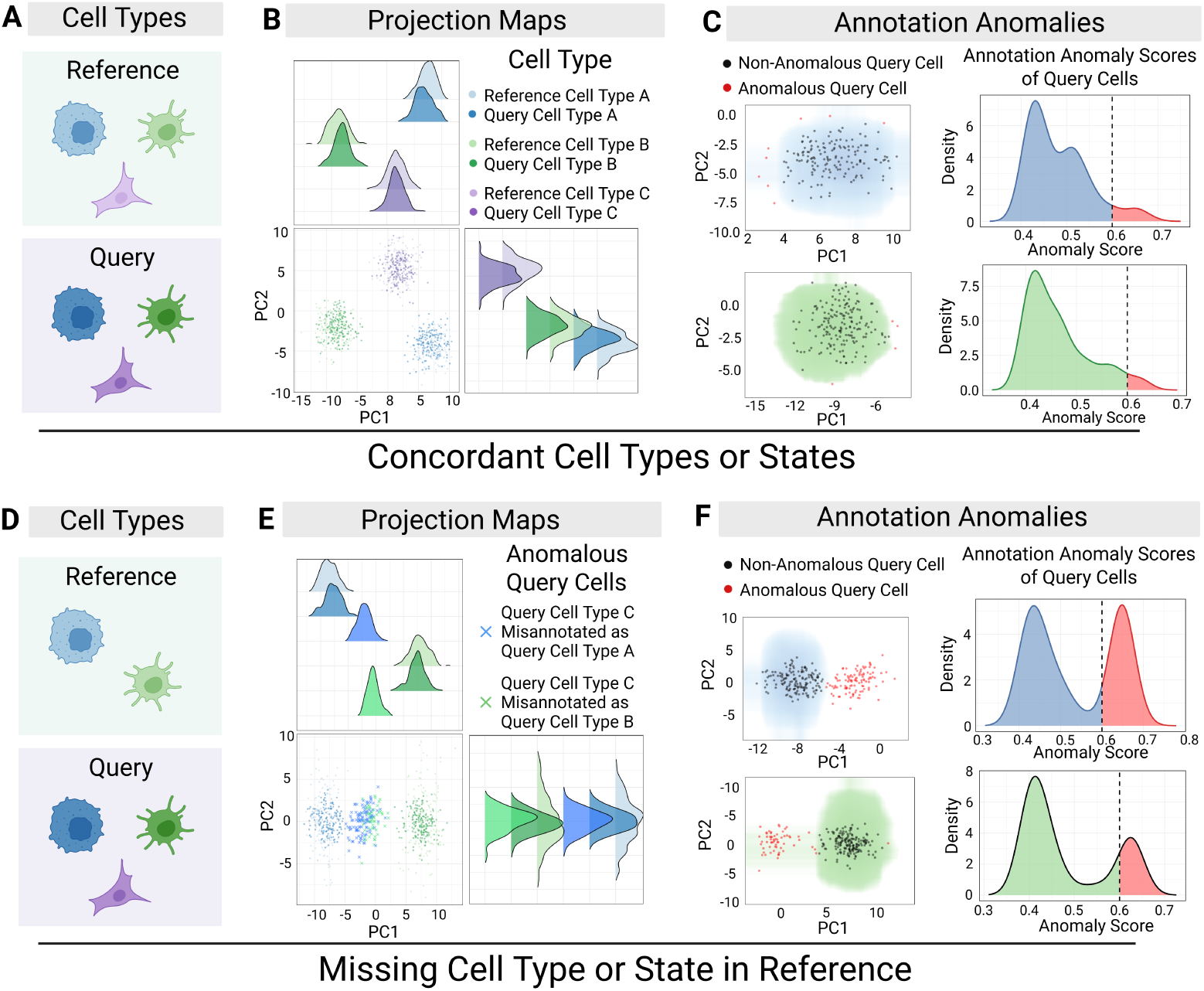
Detection of out-of-reference populations in simulated single-cell data with scDiagnostics. (**A**) Simulation setup 1: reference and query datasets are composed of the same cell types A, B, and C. (**B**) Query cells projected onto the reference embedding show concordant alignment across cell types. (**C**) Diagnostic plots show non-anomalous (black) and anomalous (red) query cells, with anomaly score distributions indicating cell populations that are well-separated within datasets and well-aligned between datasets. (**D**) Simulation setup 2: query dataset contains green cell type C not present in the reference. (**E**) Query cells of type C are misannotated as types A or B, but are flagged as an anomalous population (highlighted in red) by `scDiagnostics`. (**F**) Elevated anomaly scores successfully isolate misannotated query cells, separating them from non-anomalous reference-like populations.

In the first scenario, both datasets contained the same set of cell types A, B and C (Fig. 2A, “concordant cell types or states”). After cell type label transfer, projection plots showed precise alignment of query cells with their reference counterparts (Fig. 2B). Consequently, anomaly scores for the query cells followed a distribution nearly identical to that of the reference cells, with uniformly low scores indicating a successful integration and correct annotation (Fig. 2C).

In the second scenario, we removed cell type C from the reference dataset but retained it in the query (Fig. 2D, “missing cell type or state”). Standard reference-based annotation methods forcibly assigned these type C query cells one of the two available reference cell type labels, resulting in misclassifying these cells as either cell type A or B. `scDiagnostics` correctly identified these cells as misannotated (Fig. 2E). The projection analysis revealed that the misclassified type C cells formed a distinct cluster offset from the reference type A and B manifolds. Importantly, the anomaly detection module flagged these cells as anomalous with significantly elevated anomaly scores, as apparent from a bimodal distribution that clearly separated correctly annotated cells from the out-of-reference population (Fig. 2F).

To simulate out-of-reference populations also in real single-cell data, we next evaluated `scDiagnostics` on the Zeisel mouse brain scRNA-seq dataset [25], a well-characterized reference atlas encompassing all major cell populations in the cortex, including neurons, glia, and vascular cells. We therefore systematically withheld specific cell populations from the reference, including (i) a distinct glial lineage (astrocytes), (ii) one of two closely related neuronal subtypes, and (iii) a rare population (microglia). To further reflect a range of complexities typically encountered in real single-cell data, we introduced controlled gradients of technical and biological confounders: (i) annotation label noise (up to 40% shuffled labels), (ii) class imbalance (reducing reference populations down to 10 cells), and (iii) simulated batch effects (up to +2 mean log-count shifts). Across these realistic scenarios, anomaly detection demonstrated high areas under both the receiver operating characteristic (AUROC) and precision-recall (AUPRC) curves, alongside high sensitivity and specificity (Supplementary Results S2.1 and Supplementary Figures S3–S6).

Having established the reliability of `scDiagnostics` in controlled settings with known ground truth, we next applied the diagnostic framework to the cell type annotation of two additional real-world single-cell datasets. In the following sections, we demonstrate how the application of `scDiagnostics` to (i) single-cell RNA-seq data of COVID-19 patients, and (ii) spatial transcriptomics data of a mouse model of colitis, significantly improves the accuracy of downstream biological interpretation of the data.

### scDiagnostics facilitates the identification and characterization of a disease-associated cell state in COVID-19

To demonstrate the package’s capabilities to recover known disease-specific cell states that would otherwise be missed by standard reference-based annotation workflows, we next applied `scDiagnostics` to the single-cell peripheral blood mononuclear cell (PBMC) atlas generated by Stephenson et al. [26] (Fig. 3). We deliberately selected this dataset because severe COVID-19 infection induces a well-documented, complex transcriptomic shift in monocytes [20, 27], providing a reliable real-world positive control to validate our diagnostic framework. We partitioned the data into (i) a reference dataset comprising 24 healthy donors, and (ii) a query dataset comprising 100 COVID-19 patients of varying disease severity (Fig. 3A, left). Cell type labels were transferred from the healthy reference to the disease query using Seurat’s [16] `Azimuth` framework [17] (Fig. 3A, center). To determine the inflammatory status of all annotated cells, we integrated both datasets and calculated a Type I interferon (IFN) gene signature score as previously described by Yoshida et al. [27] (Fig. 3A, right).

**Fig. 3.**
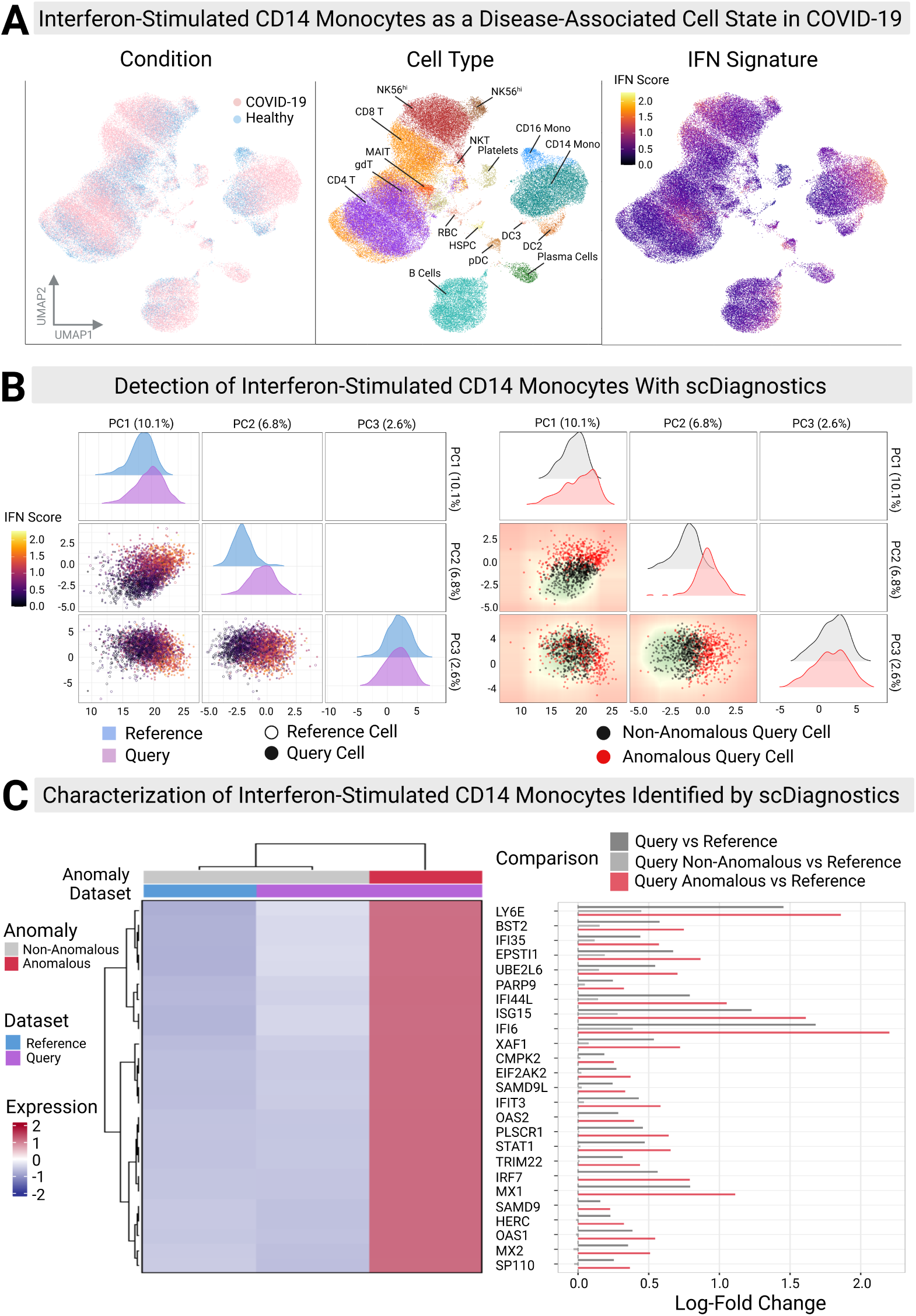
Detection and characterization of a disease-associated cell state in COVID-19 with scDiagnostics. (**A**) UMAP visualization of integrated healthy (reference) and COVID-19 (query) PBMCs colored by condition (left), cell type (center), and interferon (IFN) signature expression (right). (**B**) PCA projection reveals a deviating cell state correlated with high IFN scores (left), successfully isolating the anomalous query population (right, red points) that display a shift from the reference manifold. (**C**) Characterization of the highlighted anomalous query cells with differential expression analysis. Heatmap (left) and log-fold change analysis (right) display increased expression of specific IFN signature genes in anomalous query cells compared to the reference.

Although the global UMAP embedding indicated reasonable alignment between reference and query for most immune cell populations, `scDiagnostics` revealed a significant distributional shift within the myeloid cell class. Specifically, a subset of cells annotated as CD14^+^ monocytes in the COVID-19 query dataset showed strongly elevated anomaly scores, displaying a bimodal distribution that was significantly different from the unimodal anomaly score distribution of the reference data (Fig. 3B, right). In principal component (PC) space, these high-scoring cells were markedly shifted away from the center of the healthy monocyte manifold (Fig. 3B, left). Furthermore, overlaying the expression score of the IFN signature onto the low-dimensional projection revealed a clear concordance: cells flagged as anomalous (red points in Fig. 3B, right) displayed high IFN signature expression (orange/yellow gradient). This apparent visual correlation indicated that the identified outliers indeed correspond to the disease-associated interferon-enriched cell state.

To further characterize the anomalous CD14^+^ monocyte population, we applied pseudobulk differential expression analysis on cells stratified by anomaly status within the 25-gene IFN signature from Yoshida et al. [27]. The resulting top differentially expressed genes included many members of the IFN signature, confirming significant expression differences between the anomalous and non-anomalous query populations (Fig. 3C, heatmap). Strongest expression differences were observed for *IFI6*, *LY6E*, and *ISG15*, with log_2_-fold changes of 2.30, 1.94, and 1.72 in anomalous vs. healthy reference CD14^+^ monocytes (Fig. 3C, red bars). In contrast, non-anomalous CD14^+^ monocytes displayed markedly lower expression differences for these genes (log_2_-fold changes of 0.49, 0.53, and 0.36, respectively; Fig. 3C, light grey bars). Across all 25 IFN-response genes, expression changes were significantly higher in anomalous vs. non-anomalous CD14^+^ monocytes (mean log_2_-fold change 0.78 *±* 0.52 vs. 0.13 *±* 0.14, *p* = 2.21 *×* 10^−8^, paired *t*-test). The strongest differences were observed for *IFI6*, *LY6E*, and *ISG15* (difference in log_2_-fold change: 1.81, 1.41, and 1.35, respectively).

Critically, when treating all query CD14^+^ monocytes as a single population as resulting from the standard reference-based annotation, this clear separation was obscured, with significantly reduced log_2_-fold changes of 0.60 *±* 0.41 vs. 0.78 *±* 0.52 for anomalous CD14^+^ monocytes (Fig. 3C, dark grey bars, *p* = 2.16 *×* 10*^−^*^8^, paired *t*-test). These findings are in agreement with hyper-inflammatory monocyte states characterized by extensive IFN upregulation as previously described for SARS-CoV-2 infection [20], demonstrating that `scDiagnostics` facilitates the identification of disease-specific CD14^+^ monocytes with characteristic IFN expression profiles that the standard reference-based annotation conflated with healthy reference monocytes.

We also note that these findings can be replicated when applying `scDiagnostics` to cell type labels obtained from `SingleR` [12], `CellTypist` [13], and scArches [19] (Supplementary Results S2.2). Across all major annotation tools, `scDiagnostics` consistently identified the interferon-enriched CD14^+^ monocyte population as anomalous, demonstrating that the recovery of this disease-associated cell state is independent of the underlying annotation method. Notably, many of these transcriptomically divergent cells retained high confidence scores across multiple annotation tools, illustrating that annotation tool confidence metrics alone do not reliably capture dataset incompatibilities or disease-driven transcriptional shifts (Supplementary Results S2.2, Supplementary Figures S9–S13, and Supplementary Tables S2–S4).

### scDiagnostics facilitates the identification and characterization of a disease-associated cell state in colitis

To also demonstrate the application of `scDiagnostics` to spatially-resolved transcriptomics data, we next analyzed MERFISH imaging-based transcriptomics of a dextran sodium sulfate (DSS)-induced mouse model of colitis from Cadinu et al. [28] (Fig. 4). We compared a colitis tissue slice from Day 9 (query) against a healthy colon slice from Day 0 (reference). Similar to the COVID-19 analysis, this dataset serves as a biological positive control, as DSS-induced colitis is known to drive the emergence of spatially distinct inflammation-associated fibroblasts [28, 29]. The resulting cell type annotations showed a clear degradation of the epithelial membrane (Fig. 4A, left), and spatial localization of inflammation-associated fibroblasts (IAFs) along the inflamed epithelium (Fig. 4A, center), a cell state previously implicated in pathological remodeling in inflammatory bowel disease (IBD) [29].

**Fig. 4.**
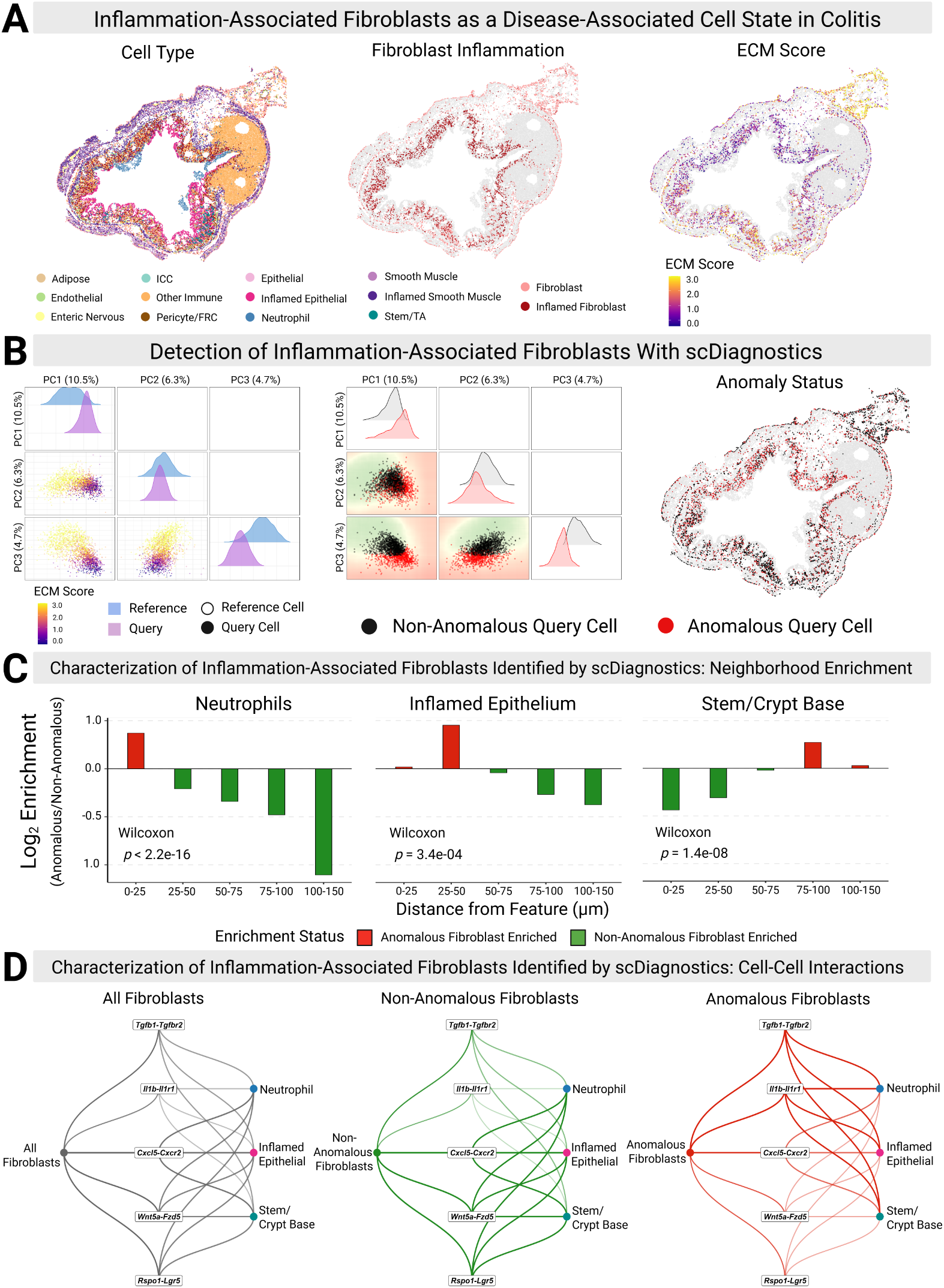
Detection and characterization of a disease-associated cell state in a mouse model of colitis with scDiagnostics. (**A**) Colon tissue slice at the onset of colitis showing cell type annotations (left), spatial segregation of normal and inflamed fibroblasts (center), and extracellular matrix (ECM) expression scores (right). (**B**) PCA projection (left)identifies the anomalous query population (center, red points) associated with high ECM scores, which correspond to inflamed fibroblasts when mapped back onto the tissue slice (right). (**C**) Neighborhood enrichment analysis. The identified anomalous fibroblasts are spatially enriched near infiltrating neutrophils and inflamed epithelium, whereas non-anomalous fibroblasts are predominantly located near the crypt base. (**D**) Cell-cell interaction analysis. Non-anomalous fibroblasts maintain homeostatic Wnt signaling with stem cells, while the flagged anomalous fibroblasts display characteristics of inflammatory signaling through increased interaction with infiltrating neutrophils and inflamed epithelium.

Diagnostic analysis identified a distinct subpopulation of query cells that deviated significantly from the homeostatic fibroblast manifold. To determine the biological identity of these anomalous cells, we calculated an extracellular matrix (ECM) homeostasis signature score using canonical ECM marker genes [30] (e.g., *Col1a2*, *Timp2*, *Sparc*, *Dpt*) (Fig. 4A, right). The anomaly scores correlated strongly with the expression scores of the ECM signature (Fig. 4B).

Taking into account the spatial context of the identified anomalous fibroblast population, we performed a neighborhood enrichment analysis to further characterize these cells and their microenvironmental niche. This revealed distinct patterns of spatial co-localization of fibroblasts with infiltrating neutrophils, inflamed epithelial cells, and crypt base stem cells (Fig. 4C). Anomalous fibroblasts were significantly closer to infiltrating neutrophils (*p <* 2.2 *×* 10*^−^*^16^, Wilcoxon rank-sum test) and inflamed epithelium (*p* = 3.4 *×* 10*^−^*^4^) compared to their non-anomalous counterparts. Visualization of enrichment across distance bins showed that infiltrating immune cells were strongly overrepresented within 0–50 *µ*m of anomalous fibroblasts (Fig. 4C, left/center). In contrast, non-anomalous fibroblasts were significantly enriched near the base of the crypt (*p* = 1.4 *×* 10*^−^*^8^), consistent with the homeostatic role of fibroblasts in maintaining the intestinal stem cell niche [31].

This diagnostic fibroblast stratification was critical for accurately resolving cell-cell communication networks, as inferred from ligand-receptor co-expression analysis (Fig. 4D). The analysis yielded a dense, non-specific network when all fibroblasts were subsumed into one cell group, conflating homeostatic and inflammatory signals (Fig. 4D, left). However, separating the population according to anomaly status revealed a substantial rewiring of the interaction network. Non-anomalous fibroblasts maintained interactions involved in Wnt signaling (e.g., *Wnt5a*-*Fzd5*, *Rspo1* -*Lgr5*) with stem cells. In contrast, anomalous fibroblasts exhibited a loss of these homeostatic signals and a gain of inflammatory interactions (e.g., *Il1b*-*Il1r1*, *Cxcl5* -*Cxcr2*) with the neutrophil infiltrate (Fig. 4D, right). These results illustrate that `scDiagnostics` can effectively decompose spatially-resolved microenvironments, separating cells actively contributing to the disease pathology from healthy cells.

To demonstrate the compatibility of scDiagnostic with popular annotation tools, we applied `scDiagnostics` to cell type labels obtained from `SingleR` [12], `Azimuth` [17], `CellTypist` [13], and scVI/scArches [19, 32] (Supplementary Results S2.3, Supplementary Figures S14–S19, and Supplementary Table S5). Across all annotation tools, `scDiagnostics` consistently identified the inflammatory fibroblast population as anomalous, demonstrating that this finding can be replicated when using automated annotation transfer for cell type annotation.

## Discussion

The `scDiagnostics` package implements a computational framework for the evaluation of cell type annotations and the identification of anomalous cell states. As single-cell datasets grow into the millions of cells, researchers and single-cell analysts increasingly rely on automated reference mapping methods; however, these methods often operate under the assumption that the query data structurally mirrors the reference. Our results from applying `scDiagnostics` to simulated data and real-world case studies demonstrate that adhering strictly to this assumption can obscure critical biological deviations. By introducing a set of simple diagnostic principles, including projection mapping, dataset alignment, and anomaly detection, into the standard cell type annotation workflow, we provide straightforward mechanisms to identify disease-associated cell states that would otherwise be forcibly integrated into healthy manifolds, addressing a critical gap in automated evaluation strategies for case-control settings [33].

Accurate cell type annotation is intrinsically linked to the challenge of distinguishing genuine biological variation from technical effects during batch correction and data integration. Several benchmarking studies have shown that many established integration methods tend to “over-correct” batch effects by flattening out true biological signals, resulting in the undesired merging of distinct cell states to satisfy topological alignment between batches and datasets [34]. Such instances of over-correction can readily be identified with `scDiagnostics`. For example, the application of `scDiagnostics` to the COVID-19 PBMC atlas from Stephenson et al. [26] demonstrated that the integration strategy correctly aligned healthy CD14^+^ monocytes between reference and query dataset, but merged the COVID19-associated interferon-stimulated CD14^+^ monocytes into a cluster with the healthy CD14^+^ monocytes from the reference dataset. The statistical metrics and diagnostic tools implemented in the `scDiagnostics` package therefore allow users to quantitatively assess whether an integration is biologically meaningful or rather forces disease-specific cell states into their healthy reference baseline (see also Supplementary Discussion S3).

A recurring difficulty for users is knowing how to interpret a flagged anomaly, and in particular how to distinguish a biologically meaningful deviation from an annotation error or a technical artifact. We therefore recommend a practical interpretation approach that combines the two scores available to the user—the `scDiagnostics` anomaly score and the confidence score reported by the annotation method—with the package’s alignment and marker-gene diagnostics (Supplementary Figure S1). Crucially, the two scores are largely complementary: a high confidence score does not preclude a high anomaly score, and it is precisely these confidently mislabelled cells that standard workflows fail to detect (Supplementary Figure S1A). Indeed, in the COVID-19 PBMC data the interferon-stimulated CD14+ monocytes retained high confidence across multiple annotation tools while being clearly flagged as anomalous, underscoring that confidence metrics alone do not capture dataset incompatibilities or disease-driven transcriptional shifts. Once a cell is flagged as anomalous, the user can interrogate potential causes—technical artifact, biological drift within a type, a missing reference state, or annotation/model failure—by examining the corresponding diagnostic signature (Supplementary Figure S1B). We emphasize that these categories are adjudicated by complementary evidence rather than by the anomaly score in isolation: marker-gene alignment and quality-control associations flag technical confounders, reproducibility of a coherent signature across annotation tools points to genuine biological drift, and the absence of a matched reference population identifies out-of-reference states. This framework operationalizes the distinction between novel or shifted cell types or states and annotation failures, and provides a transparent, human-in-the-loop approach for integrating `scDiagnostics` into routine annotation workflows.

The development of `scDiagnostics` comes at a time of emerging single-cell analysis paradigm shifts driven by single-cell foundation models. These large-scale models trained on massive cell censuses promise a universal interface to many single-cell analysis and annotation tasks through representation learning [11, 35–37]. Although these approaches have the potential to eventually replace specialist models for individual single-cell analysis tasks, recent benchmark studies suggest that complex deep-learning models do not yet consistently outperform simpler linear baselines or standard statistical approaches for specific tasks [38], especially in zero-shot settings where the model encounters unseen biological states [39]. `scDiagnostics` relies on interpretable, statistical foundations offering a transparent and lightweight approach also for complementing cell type annotation with foundation models, in particular in situations where these models hallucinate a match for an unknown cell state.

Similarly, the recent application of large language models (LLMs) for cell type annotation offers new opportunities for automation and streamlining of manual marker-based cell type annotation of empirically-derived cell clusters [40]. However, LLMs lack an inherent sense of quantitative uncertainty regarding the underlying marker gene expression data. We envision `scDiagnostics` acting as a quantitative “guardrail” for such AI-driven workflows. If an LLM or foundation model confidently assigns a cell type label to a cell, but `scDiagnostics` flags it as highly anomalous, this discrepancy can trigger a “human-in-the-loop” review, preventing the propagation of high-confidence misannotations in the cell type annotation step to downstream analysis and biological interpretation.

Although `scDiagnostics` has been developed with scRNA-seq data in mind, we have shown that many of the implemented diagnostic principles can be straightforwardly applied to spatially-resolved single-cell transcriptomics data. The application to the MERFISH colitis timecourse from Cadinu et al. [28] demonstrated that `scDiagnostics` can play an important role by identifying cells where the expression profile conflicts with the spatial context – for example, fibroblasts displaying an inflammatory profile due to proximity to infiltrating neutrophils. This spatially-aware anomaly detection is therefore essential for distinguishing cells actively driving tissue pathology from healthy reference-like cells. On the other hand, several extensions of `scDiagnostics` are warranted for data obtained from (i) sequencing-based approaches with supracellular resolution, which require deconvolution of cell type mixtures [41], and (ii) imaging-based approaches with subcellular resolution that rely on cell segmentation [42]. For sequencing-based spatial transcriptomics, `scDiagnostics` can be naturally extended to evaluate deconvolution quality by adapting the existing functions for comparing query and reference datasets to assess the compatibility between the spatial data and scRNA-seq references used for deconvolution. For imaging-based spatial transcriptomics, recent advances in joint segmentation and annotation aim to refine cell boundaries using gene expression [42, 43], creating new opportunities for diagnostic evaluation. For example, diagnostics can be developed to assess the consistency between cell morphology and cell type annotation, identifying cases where the assigned cell type is inconsistent with the size, shape, or spatial context of the cell.

We also note that although this work is focused on transcriptomics data, the diagnostic framework presented here can be extended and applied also to other single-cell data modalities. As single-cell multi-modal data is becoming increasingly commonplace, complementary aspects and discrepancies between modalities—such as disagreements between scRNA-seq data and surface protein expression in CITE-seq [44] data—are typically ignored or only insufficiently investigated [45]. Additional avenues for future development of the package could therefore focus on multi-modal anomaly detection, enabling researchers to identify cells where measurements of different modalities are not in agreement with the central dogma due to biological processes such as post-transcriptional regulation, or technical factors such as non-specific antibody binding and compromised mRNA integrity. By formalizing the search for outliers and anomalies, multi-modal diagnostics could thus transform the analysis of these cells from a nuisance into a primary source of biological discovery.

## Conclusions

The `scDiagnostics` package provides a critical set of diagnostic tools for assessing cell type annotation in single-cell transcriptomics studies, addressing the unmet need for systematic, method-agnostic evaluation of automated cell type annotation methods. By integrating projection-based visualization, quantitative dataset and marker alignment metrics, and detection of annotation anomalies, `scDiagnostics` facilitates the identification and characterization of misannotated, out-of-reference, novel or disease-associated cell types or states that would otherwise remain obscured in standard workflows. Based on carefully simulated and diverse real-world datasets, we demonstrated that this diagnostic approach significantly improves downstream analysis and biological interpretation, recovering disease signal in differential expression analysis, and revealing functional rewiring in cell–cell communication networks. As single-cell datasets continue to grow in scale and complexity, and as foundation models increasingly automate annotation, tools that systematically evaluate annotation reliability are essential for maintaining rigor and enabling reproducible biological discovery. We therefore hope that `scDiagnostics` provides single-cell analysts with a useful toolkit to routinely interrogate and refine their cell type annotations, thereby enhancing the reliability and interpretability of biological conclusions.

## Methods

### Implementation

The `scDiagnostics` package is implemented in R [46] and is part of the Bioconductor [47] ecosystem. Designed for broad interoperability with major single-cell analysis workflows, `scDiagnostics` accepts `SingleCellExperiment` objects [4] as input. Notably, the package natively supports on-disk, HDF5-backed data structures via the `DelayedArray` framework [48], enabling memory-efficient processing of out-of-core, atlas-scale datasets. This widely-used data format allows users to assess annotations generated by common R-based tools, including `SingleR` [12] and Seurat’s [16] `Azimuth` [17], as Seurat objects are readily converted to the required input format. Integration with Python-based tools such as `CellTypist` [13] and `scArches` [19] is facilitated by conversion utilities such as `reticulate` [49] and `zellkonverter` [50], bridging the gap between the `AnnData` format [51] and R-based data infrastructure.

### Exploratory data analysis of cell type annotations

The `scDiagnostics` package offers visual diagnostics of cell type annotation quality by mapping the query dataset onto a low-dimensional representation of the reference dataset. This projection approach facilitates direct comparison of cell type distributions, revealing potential compositional differences or annotation inconsistencies. Visualization strategies include: principal component analysis (PCA) projections that capture overall variation across genes [52]; multidimensional scaling (MDS) plots emphasizing similarity relationships [53]; discriminant analysis projections highlighting genes with the strongest differentiating power [54]; and sliced inverse regression (SIR) functionality to identify directions optimally separating cell types in high-dimensional space [55]. Additional functions assess how common quality control (QC) metrics influence annotation confidence scores, enabling identification of technical factors compromising accuracy.

### Evaluation of dataset and marker gene alignment

The `scDiagnostics` package evaluates dataset alignment through statistical approaches that project query data into the PCA-derived reference subspace to assess distributional congruence. Alignment metrics computed within this shared space include: Spearman correlation analysis to assess cell-cell relationships; optimal transport-based Wasserstein distance calculations [56, 57], Bhattacharyya coefficients [58], and Hellinger distances [59] to quantify the magnitude of structural divergence and flag potential integration failures; multivariate hypothesis testing through Cramér’s test to assess cumulative distribution differences [60, 61]; and permutation-based Hotelling’s *T* ^2^ tests to evaluate multivariate mean differences [62]. For marker gene alignment assessment, the package examines agreement of overall and cell-type specific marker gene expression profiles using transformations, including Z-score standardization [63] and quantile rank alignment [64], to ensure fair comparisons. Additionally, the package computes highly variable gene (HVG) overlap coefficients [34] and employs random forest variable importance measures to identify genes most effective in distinguishing cell types [65].

### Detection and analysis of annotation anomalies

The `scDiagnostics` package identifies anomalous cell type annotations through a combination of PCA projection with outlier detection using the isolation forest algorithm [66], as well as a complementary geometric baseline evaluating PCA reconstruction error. This involves projecting query data into the PCA-derived coordinate system of the reference, enabling detection of outlying query cells deviating from expression patterns observed in the reference. Isolation forests identify outliers by exploiting their characteristic property of requiring fewer random partitions for isolation, and have recently been adapted and validated for high-dimensional omics datasets, where they recover clinically meaningful outliers and biomarker patterns [67–69]. Within `scDiagnostics`, users can run isolation forests either on principal component (PC) scores or directly on a restricted set of highly variable genes, and can choose between cell-type-specific anomaly detection, where models are trained on individual cell populations, and global detection across all cell types simultaneously. Anomaly thresholds can be specified either as a fixed absolute cutoff on the isolation-forest score, or in a data-adaptive manner based on the median and MAD of reference scores. Supplementary Results S2.1 and Supplementary Figures S7–S8 examine the impact of these choices on sensitivity and specificity.

### Simulation framework

We simulated synthetic scRNA-seq datasets using the splatter framework [70] to generate a balanced two-batch dataset (*n* = 500 cells per batch) with three distinct cell types (A, B, C) of equal abundance. Transcriptomic profiles used differential expression probabilities of 0.1, 0.2, and 0.2, respectively, with outlier gene probabilities set to zero to ensure anomalies arise from structural differences rather than stochastic noise.

To illustrate shortcomings of reference-based annotation, we introduced a composition mismatch. The reference dataset retained only cell types A and B, while the query retained all three types. Following log-normalization and dimensionality reduction (PCA) via scater [71], the query was annotated with `SingleR` [12], causing the unrepresented type C cells to be misclassified into reference categories A and B.

We applied `scDiagnostics` to detect these misclassifications. Query data was projected onto the reference PCA manifold using `detectAnomaly` with an isolation forest (1,000 trees) trained on the reference. Diagnostic performance was assessed by comparing score distributions of correctly annotated versus misclassified cells. Guided by a data-adaptive approach based on the median and MAD of the reference scores, we applied an anomaly score threshold of 0.6 to identify high-confidence anomalies. Visualization was performed using the `plotCellTypePCA` and `detectAnomaly` functions.

To evaluate diagnostic performance under complex confounders frequently encountered in real single-cell data, we utilized the well-characterized Zeisel mouse brain scRNA-seq atlas [25]. Following log-normalization, the dataset was randomly partitioned into a 70% reference and 30% query subset. Out-of-distribution scenarios were constructed by withholding specific cell populations to represent diverse mapping challenges: a distinct glial lineage (astrocytes), one of two closely related neuronal subtypes (pyramidal SS), and a rare population (microglia). The query dataset was annotated using `SingleR` [12], which forcibly mapped these unrepresented populations to the most transcriptionally similar available reference categories. To assess performance under suboptimal mapping conditions, we introduced controlled gradients of technical and biological confounders: annotation label noise (randomly shuffling 10% to 40% of reference labels), progressive class imbalance (downsampling target reference populations from 300 to 10 cells), and simulated batch effects (applying mean shifts of 0 to +2 log-normalized counts to a random 20% of query genes). Anomaly detection was performed using the `detectAnomaly` and `calculateReconstructionError` functions, and diagnostic accuracy was quantified against ground-truth labels using the area under the receiver operating characteristic (AUROC) and precision-recall (AUPRC) curves, alongside sensitivity and specificity.

### Application to COVID-19 PBMC scRNA-seq data

#### Data processing and reference construction

We analyzed scRNA-seq data from the multi-center PBMC atlas generated by Stephenson et al. [26] to assess the capacity of `scDiagnostics` to detect misclassifications resulting from disease-driven transcriptional shifts. To ensure a balanced experimental design, we performed stratified subsampling within each sequencing site, retaining up to 1,000 cells per healthy sample (reference) and exactly 500 cells per COVID-19 sample (query). This minimized site-specific batch effects, ensuring that primary disease-reference comparisons were not confounded by technical variation.

QC was performed using `scater` [71], removing cells with fewer than 1,000 UMIs and genes expressed in fewer than 10 cells. Counts were log-normalized, and HVGs identified using scran [72]. To ensure that the feature space captured both homeostatic and disease-driven variance, we computed the union of HVGs from both reference and query datasets prior to dimensionality reduction using PCA. Following QC and filtering, measurements for 18,472 genes were retained for further analysis across 23,201 cells in the healthy reference dataset and 48,148 cells in the COVID-19 query dataset.

#### Cell type annotation

To demonstrate compatibility with major annotation tools, the COVID-19 query dataset was annotated based on the healthy reference using four different methods (Supplementary Results S2.2). For the primary application presented here, cell type annotation was performed with `Azimuth` [17], using Seurat v3 normalization (SCT) [16] and UMAP embedding on the reference, mapping query cells onto the reference embedding. Additional analyses using `SingleR` [12], `CellTypist` [13], and `scVI/scArches` [19, 32] are available in the manuscript repository. For annotation with scVI/scArches [19, 32], we trained a variational autoencoder with `scVI` [32] on the reference data to learn a shared latent representation, using transfer learning to map query cells and assigning labels via *k*-nearest neighbors (*k* = 50).

#### Dataset integration and visualization

To visualize the integrated reference and query data, we derived UMAP embeddings from the shared latent representation computed by the trained `scVI`/`scArches` model. Cells were colored by sample condition, cell type annotation (transferred via `Azimuth` [17]), and aggregated expression level of the IFN-response signature.

#### Anomaly detection and IFN signature expression

Based on previous findings by Dann et al. [20], we investigated the CD14^+^ monocyte population for disease-associated variation. We applied the detectAnomaly function to project COVID-19 CD14^+^ monocytes onto the PCA manifold defined by healthy CD14^+^ monocytes (genome-wide), using the isolation forest algorithm to calculate anomaly scores. Anomalous cells were identified using an anomaly score threshold of 0.5, derived from a data-adaptive approach based on the median and MAD of the healthy reference anomaly scores. To determine biological relevance, we calculated an IFN-response signature score using 25 canonical IFN-response genes [27] and visualized the score across the first three PCs using the `projectPCA` function.

#### Characterization of IFN-driven anomalies

To characterize molecular drivers of the detected anomalies, we applied the `calculateGeneShifts` function to the 25-gene Yoshida et al. IFN signature [27]. This analysis performed anomaly detection on the IFN signature genes only, enabling focused comparison of pseudo-bulk expression profiles between healthy reference and anomaly-stratified query populations. Log_2_-fold changes were calculated for three pairwise comparisons: (i) anomalous query vs. reference, (ii) non-anomalous query vs. reference, and (iii) all query cells vs. reference. Paired *t*-tests were used to assess the statistical significance of fold-change differences.

### Application to MERFISH spatial transcriptomics data

#### Data processing and reference construction

To demonstrate the application of `scDiagnostics` to spatially-resolved single-cell data, we analyzed MERFISH data from Cadinu et al. [28], profiling a mouse model of DSS-induced colitis over a 21-day timecourse (peak colitis at Day 9). We compared spatially-resolved expression of mouse colon tissue slices between two timepoints: Day 0 (healthy baseline, Mouse ID 082421-D0-m6, Slice 2) and Day 9 (induced colitis, Mouse ID 062221-D9-m3, Slice 2).

Data was obtained from the `MerfishData` Bioconductor package [73] as integrated analysis-ready `SpatialExperiment` [74] objects linking expression measurements and cell type annotations with spatial coordinates. Given that `scDiagnostics` utilizes Bioconductor’s `SingleCellExperiment` container [4], it is natively compatible with `SpatialExperiment` objects [74], enabling seamless integration of spatial context into the anomaly detection process.

QC was performed using scater [71], removing outlier cells based on library size, feature detection, and mitochondrial percentage using adaptive thresholds. Cells annotated as inflammation-associated (inflammation-associated epithelial cells [IAE], inflammation-associated fibroblasts [IAF], and inflammation-associated smooth muscle cells [IASMC]) were removed from the Day 0 reference as artifacts of the manual marker-based annotation performed by the authors. Log-normalization was performed using scater [71]. Following QC, measurements for 943 genes were retained across 27,140 cells in the Day 0 reference dataset and 29,040 cells in the Day 9 query dataset. PCA was performed on the reference using scater [71].

#### Cell type annotation

For diagnostic evaluation, we used the ground-truth annotations provided by the authors. To create a scenario where disease-specific states are obscured by coarse-grained classification, we aggregated all fibroblast subtypes in the Day 9 dataset into an overall fibroblast lineage class. As for the COVID-19 analysis, cell type annotation was also performed with `SingleR` [12], `Azimuth` [17], `CellTypist` [13], and scVI/scArches [19, 32] to demonstrate compatibility (Supplementary Results S2.3).

#### Anomaly detection and ECM signature expression

We applied the `projectPCA` function to visualize query fibroblasts within the PC space defined by Day 0 reference fibroblasts. As previously described by Cadinu et al. [28], we calculated an ECM homeostasis signature score using five canonical matrisome genes [30]: *Col1a2*, *Timp2*, *Col6a1*, *Sparc*, and *Dpt*. The signature score was computed as the mean log-normalized expression, winsorized at 1.5 to reduce the influence of extreme values, and scaled by a factor of 2. We then applied the `detectAnomaly` function to the fibroblast lineage, projecting query fibroblasts onto the PCA manifold defined by Day 0 reference fibroblasts and calculating anomaly scores using the isolation forest algorithm. Anomalies were flagged using a threshold of 0.5, guided by the median and MAD of the reference scores.

#### Neighborhood enrichment analysis

To evaluate the spatial organization of anomalous vs. non-anomalous fibroblasts, we calculated Euclidean distances from each query fibroblast to specific target cell populations: neutrophils (acute inflammation), inflamed epithelium (tissue damage), and stem/crypt base cells (homeostatic niche). Distances were computed using the `FNN` package [75] (*k* = 1). We quantified spatial enrichment by binning distances into intervals (0–25, 25–50, 50–75, 75–100, 100–150, and *>* 150 *µm*) and calculating the log_2_ fold change of cell proportions for anomalous vs. non-anomalous fibroblasts. Statistical significance was assessed using two-sided Wilcoxon rank-sum tests.

#### Cell-cell interaction analysis

To characterize interaction networks, we modeled ligand-receptor interaction potential as the product of mean ligand expression in sender fibroblasts and mean receptor expression in receiver cell types. We used a curated set of six interaction pairs determined by Cadinu et al. [28]: *Wnt5a*-*Fzd5*, *Rspo1* -*Lgr5*, *Il1b*-*Il1r1*, *Il11* -*Il11ra*, *Tgfb1* -*Tgfbr2*, and *Cxcl5* -*Cxcr2*. Scores were independently computed for three sender categories: anomalous fibroblasts, non-anomalous fibroblast, and all fibroblast combined (baseline). Scores were normalized relative to the maximum value for each ligand-receptor pair. Interaction networks were visualized using ggraph [76] and tidygraph [77], with edge width and opacity proportional to interaction strength.

## Data availability

The COVID-19 PBMC data was obtained from the multi-center PBMC atlas generated by Stephenson et al. [26], which is available from the CELLxGENE portal [78] (collection ID: ddfad306-714d-4cc0-9985-d9072820c530). The MERFISH mouse colitis dataset generated by Cadinu et al. [28] was obtained from the `MerfishData` Bioconductor package [73]. This package provides analysis-ready `SpatialExperiment` [74] objects linking expression measurements and cell type annotations with spatial coordinates.

## Code availability

The code for data processing, analysis, and figure generation is available in the manuscript repository at https://github.com/ccb-hms/scDiagnosticsManuscript. A companion website with tutorials and organized workflows for replicating results is available at https://ccb-hms.github.io/scDiagnosticsManuscript/. The repository includes: (i) scripts to reproduce the results presented in this manuscript; (ii) comparative annotation workflows for four major cell type annotation methods (`SingleR` [12], `Azimuth` [17], `CellTypist` [13], and scArches [19]); and (iii) user-friendly tutorials demonstrating key functionality of `scDiagnostics` on real datasets. The `scDiagnostics` R package is available from Bioconductor at https://bioconductor.org/packages/scDiagnostics. All diagnostics are also available in an interactive application to facilitate access for applied users and bench scientists at https://ccb.connect.hms.harvard.edu/scDiagnosticsApp/.

## Key Points

- `scDiagnostics` provides a systematic, method-agnostic framework to evaluate cell type annotations in single-cell transcriptomics data, addressing a major gap in existing analysis workflows where label transfer is often performed without rigorous quality assessment.
- The package combines three complementary diagnostics: reference-based projections, quantitative reference–query alignment metrics (global and marker-gene level), and anomaly/outlier detection, to flag ambiguous or out-of-reference cells that standard confidence scores can miss
- Using controlled simulations, `scDiagnostics` reliably detects forced misannotations arising when query cell states are absent from the reference, separating correctly annotated cells from out-of-reference populations via elevated anomaly scores. Quantitative benchmarking on a mouse brain atlas demonstrates high sensitivity and specificity across realistic sources of simulated noise, class imbalance, and batch effects.
- In COVID-19 PBMCs, `scDiagnostics` successfully recovers an interferon-stimulated CD14^+^ monocyte state that is conflated with healthy monocytes by reference mapping, restoring disease signal that would otherwise be diluted in downstream differential expression analyses.
- In MERFISH colitis data, `scDiagnostics` identifies inflammation-associated fibroblasts with spatially distinct niches and rewired ligand–receptor interactions, demonstrating utility beyond scRNA-seq and highlighting how diagnostic stratification can improve biological interpretation.

## Author Biographies

**Anthony Christidis** is a Computational Scientist at the Center for Computational Biomedicine, Harvard Medical School. His research focuses on the application of statistical and computational methods to single-cell multi-modal data. **Andrew Ghazi** is a Senior Computational Biologist at the Center for Computational Biomedicine, Harvard Medical School. His research focuses on probabilistic modeling of single-cell multi-modal data. **Smriti Chawla** is a Postdoctoral Research Fellow at the Center for Computational Biomedicine, Harvard Medical School. Her research focuses on statistical and computational methods for single-cell data analysis. **Nitesh Turaga** is the Director of Research Informatics at Tempus AI. He was previously a member of the Bioconductor core team and the Galaxy core team. He leads the implementation of critical software tools for multi-modal cancer data analysis. **Robert Gentleman** is a Principal Research Scientist at the Dana Farber Cancer Institute. He is one of the creators of the R programming language and one of the founders of the Bioconductor project. His research focuses on scientific computing, data visualization, genomics, genetics, machine learning, and the application of statistical and computational methods to studying human disease. **Ludwig Geistlinger** is the Director of Computational Biology at the Center for Computational Biomedicine, Harvard Medical School. He is a member of Bioconductor’s Technical Advisory Board. His research focuses on applications of statistical and computational methods in single-cell and spatial omics data analysis, gene set and network enrichment analysis, copy number variation analysis, human microbiome analysis, and multi-omic analysis in cancer genomics.

## Acknowledgements

We wish to thank our colleagues at the Center for Computational Biomedicine for their feedback on the project, and the Bioconductor Core Team for peer-review of the implementation of the `scDiagnostics` R/Bioconductor package.

## Supplementary Data

Supplementary data are available at *Briefings in Bioinformatics* online.

## References

[1] Macosko, E. Z. et al. Highly parallel genome-wide expression profiling of individual cells using nanoliter droplets. Cell 161, 1202–14 (2015).

[2] Zheng, G. X. et al. Massively parallel digital transcriptional profiling of single cells. Nat Commun 8, 14049 (2017).

[3] Luecken, M. D. & Theis, F. J. Current best practices in single-cell RNA-seq analysis: a tutorial. Mol Syst Biol 15, e8746 (2019).

[4] Amezquita, R. A. et al. Orchestrating single-cell analysis with Bioconductor. Nat Methods 17, 137–45 (2020).

[5] Lähnemann, D., et al. Eleven grand challenges in single-cell data science. Genome Biol 21, 31 (2020).

[6] Luecken, M. D. et al. Defining and benchmarking open problems in single-cell analysis. Nat Biotechnol 43, 1035–40 (2025).

[7] Clarke, Z. A. et al. Tutorial: guidelines for annotating single-cell transcriptomic maps using automated and manual methods. Nat Protoc 16, 2749–64 (2021).

[8] Pullin, J. M. & McCarthy, D. J. A comparison of marker gene selection methods for single-cell RNA sequencing data. Genome Biol 25, 56 (2024).

[9] Hu, C. et al. CellMarker 2.0: an updated database of manually curated cell markers in human/mouse and web tools based on scRNA-seq data. Nucleic Acids Res 51, D870–6 (2023).

[10] Abdelaal, T. et al. A comparison of automatic cell identification methods for single-cell RNA sequencing data. Genome Biol 20, 194 (2019).

[11] Rood, J. E. et al. The Human Cell Atlas from a cell census to a unified foundation model. Nature 637, 1065–71 (2025).

[12] Aran, D. et al. Reference-based analysis of lung single-cell sequencing reveals a transitional profibrotic macrophage. Nat Immunol 20, 163–72 (2019).

[13] Domínguez, C. C., et al. Cross-tissue immune cell analysis reveals tissue-specific features in humans. Science 376, eabl5197 (2022).

[14] de Kanter, J. K., Lijnzaad, P., Candelli, T., Margaritis, T. & Holstege, F. C. P. CHETAH: a selective, hierarchical cell type identification method for single-cell RNA sequencing. Nucleic Acids Res 47, e95 (2019).

[15] Kiselev, V. Y., Yiu, A. & Hemberg, M. scmap: projection of single-cell RNA-seq data across data sets. Nat Methods 15, 359–62 (2018).

[16] Stuart, T. et al. Comprehensive integration of single-cell data. Cell 177, 1888–902.e21 (2019).

[17] Butler, A., et al. Azimuth: a Shiny app demonstrating a query-reference mapping algorithm for single-cell data (2023). URL https://github.com/satijalab/Azimuth. R package version 0.5.0.

[18] Shao, X. et al. scDeepSort: a pre-trained cell-type annotation method for single-cell transcriptomics using deep learning with a weighted graph neural network. Nucleic Acids Res 49, e122 (2021).

[19] Lotfollahi, M. et al. Mapping single-cell data to reference atlases by transfer learning. Nat Biotechnol 40, 121–30 (2022).

[20] Dann, E. et al. Precise identification of cell states altered in disease using healthy single-cell references. Nat Genet 55, 1998–2008 (2023).

[21] Xu, J., Liao, K., Yang, X., Wu, C. & Wu, W. Using single-cell sequencing technology to detect circulating tumor cells in solid tumors. Mol Cancer 20, 104 (2021).

[22] Kotliar, D. et al. Reproducible single-cell annotation of programs underlying T cell subsets, activation states and functions. Nat Methods 22, 1964–80 (2025).

[23] Pijuan-Sala, B. et al. A single-cell molecular map of mouse gastrulation and early organogenesis. Nature 566, 490–5 (2019).

[24] Chari, T. & Pachter, L. The specious art of single-cell genomics. PLoS Comput Biol 19, e1011288 (2023).

[25] Zeisel, A. et al. Brain structure. cell types in the mouse cortex and hippocampus revealed by single-cell rna-seq. Science 347, 1138–1142 (2015).

[26] Stephenson, E. et al. Single-cell multi-omics analysis of the immune response in COVID-19. Nat Med 27, 904–16 (2021).

[27] Yoshida, M. et al. Local and systemic responses to SARS-CoV-2 infection in children and adults. Nature 602, 321–7 (2022).

[28] Cadinu, P. et al. Charting the cellular biogeography in colitis reveals fibroblast trajectories and coordinated spatial remodeling. Cell 187, 2010–28 (2024).

[29] Kinchen, J. et al. Structural remodeling of the human colonic mesenchyme in inflammatory bowel disease. Cell 175, 372–86 (2018).

[30] Naba, A. et al. The matrisome: in silico definition and in vivo characterization by proteomics of normal and tumor extracellular matrices. Mol Cell Proteom 11, M111.014647 (2012).

[31] Shoshkes-Carmel, M. et al. Subepithelial telocytes are an important source of Wnts that supports intestinal crypts. Nature 557, 242–6 (2018).

[32] Lopez, R., Regier, J., Cole, M. B., Jordan, M. I. & Yosef, N. Deep generative modeling for single-cell transcriptomics. Nat Methods 15, 1053–58 (2018).

33. Sikkema, L., et al. Automated evaluation of single-cell reference atlas mappings enables the identification of disease-associated cell states. bioRxiv (2025). URL 10.1101/2025.05.23.655749.

[34] Luecken, M. D. et al. Benchmarking atlas-level data integration in single-cell genomics. Nat Methods 19, 41–50 (2022).

[35] Cui, H. et al. scGPT: toward building a foundation model for single-cell multi-omics using generative AI. Nat Methods 21, 1470–80 (2024).

[36] Heimberg, G. et al. A cell atlas foundation model for scalable search of similar human cells. Nature 638, 1085–94 (2025).

[37] Williams, S. R. et al. Accelerating scRNA-seq analysis: automated cell type annotation using representation learning and vector search. bioRxiv (2025). URL 10.1101/2025.07.08.663651.

[38] Ahlmann-Eltze, C., Huber, W. & Anders, S. Deep-learning-based gene perturbation effect prediction does not yet outperform simple linear baselines. Nat Methods 22, 1657–61 (2025).

[39] Kedzierska, K. Z., Crawford, L., Amini, A. P. & Lu, A. X. Zero-shot evaluation reveals limitations of single-cell foundation models. Genome Biol 26, 101 (2025).

[40] Hou, W. & Ji, Z. Assessing GPT-4 for cell type annotation in single-cell RNA-seq analysis. Nat Methods 21, 1462–65 (2024).

[41] Cable, D. M. et al. Robust decomposition of cell type mixtures in spatial transcriptomics. Nat Biotechnol 40, 517–26 (2022).

[42] Littman, R. et al. Joint cell segmentation and cell type annotation for spatial transcriptomics. Mol Syst Biol 17, e10108 (2021).

[43] Jin, K. et al. Bering: joint cell segmentation and annotation for spatial transcriptomics with transferred graph embeddings. Nat Commun 16, 6618 (2025).

[44] Hao, Y. et al. Integrated analysis of multimodal single-cell data. Cell 184, 3573–87 (2021).

[45] Heumos, L. et al. Best practices for single-cell analysis across modalities. Nat Rev Genet 24, 550–72 (2023).

[46] R Core Team. R: a language and environment for statistical computing. R Foundation for Statistical Computing, Vienna, Austria (2024). URL https://www.R-project.org/.

[47] Gentleman, R. C. et al. Bioconductor: open software development for computational biology and bioinformatics. Genome Biol 5, R80 (2004).

[48] Pagès, H., Hickey, P. & Lun, A. DelayedArray: A unified framework for working transparently with on-disk and in-memory array-like datasets (2024). URL https://bioconductor.org/packages/DelayedArray. R package version 0.30.1.

[49] Ushey, K., Allaire, J. J. & Tang, Y. reticulate: interface to Python (2025). URL https://CRAN.R-project.org/package=reticulate. R package version 1.42.0.

[50] Zappia, L. & Lun, A. zellkonverter: conversion between scRNA-seq objects (2024). URL https://bioconductor.org/packages/zellkonverter. R package version 1.16.0.

[51] Virshup, I., Rybakov, S., Theis, F. J., Angerer, P. & Wolf, F. A. anndata: access and store annotated data matrices. J. Open Source Softw 9, 4371 (2024).

[52] Ringńer, M. What is principal component analysis? Nat Biotechnol 26, 303–4 (2008).

[53] Borg, I. & Groenen, P. Modern Multidimensional Scaling: theory and applications (Springer, 2005).

[54] Izenman, A. J. Modern multivariate statistical techniques (Springer, 2008).

[55] Woollard, M. et al. Weighted sliced inverse regression for scalable supervised dimensionality reduction of spatial transcriptomics data. bioRxiv (2025). URL 10.1101/2025.07.08.663651.

[56] Peyŕe, G. & Cuturi, M. Computational optimal transport: with applications to data science. Found Trends Mach Learn 11, 355–607 (2019).

[57] Bunne, C., Schiebinger, G., Krause, A., Regev, A. & Cuturi, M. Optimal transport for single-cell and spatial omics. Nat Rev Methods Primers 4, 58 (2024).

[58] Bhattacharyya, A. On a measure of divergence between two statistical populations defined by their probability distribution. Bulletin of the Calcutta Mathematical Society 35, 99–110 (1943).

[59] Pardo, L. Statistical inference based on divergence measures (Chapman and Hall/CRC, 2018).

[60] Baringhaus, L. & Franz, C. On a new multivariate two-sample test. J Multivar Anal 88, 190–206 (2004).

[61] Baringhaus, L. & Franz, C. Rigid motion invariant two-sample tests. Stat Sin 1333–61 (2010).

[62] Mardia, K. V., Kent, J. T. & Taylor, C. C. Multivariate analysis (John Wiley & Sons, 2024).

[63] Cheadle, C., Cho-Chung, Y. S., Becker, K. G. & Vawter, M. P. Application of z-score transformation to Affymetrix data. Appl Bioinform 2, 209–17 (2003).

[64] Bolstad, B. M., Irizarry, R. A., Åstrand, M. & Speed, T. P. A comparison of normalization methods for high density oligonucleotide array data based on variance and bias. Bioinformatics 19, 185–93 (2003).

[65] Breiman, L. Random forests. Mach Learn 45, 5–32 (2001).

[66] Liu, F. T., Ting, K. M. & Zhou, Z. H. Isolation forest. Proceedings of the Eighth IEEE International Conference on Data Mining 413–22 (2008).

[67] Buck, L. et al. Anomaly detection in mixed high-dimensional molecular data. Bioinformatics 39, btad501 (2023).

[68] Dai, Y., Wu, D., Carroll, I., Zou, F. & Zou, B. High-dimensional biomarker identification for interpretable disease prediction via machine learning models. Bioinformatics 41, btaf266 (2025).

[69] Peleg, O., Raytan, M. & Borenstein, E. Kadaif: an anomaly detection method for complex microbiome data. Bioinformatics 41, btaf520 (2025).

[70] Zappia, L., Phipson, B. & Oshlack, A. Splatter: simulation of single-cell RNA sequencing data. Genome Biol 18, 174 (2017).

[71] McCarthy, D. J., Campbell, K. R., Lun, A. T. L. & Willis, Q. F. Scater: pre-processing, quality control, normalisation and visualisation of single-cell RNA-seq data in R. Bioinformatics 33, 1179–86 (2017).

[72] Lun, A. T. L., McCarthy, D. J. & Marioni, J. C. A step-by-step workflow for low-level analysis of single-cell RNA-seq data with Bioconductor. F1000Res 5, 2122 (2016).

[73] Geistlinger, L., Moffitt, J. & Gentleman, R. MerfishData: Collection of public MER-FISH datasets (2024). URL https://bioconductor.org/packages/MerfishData. R package version 1.8.0.

[74] Righelli, D. et al. SpatialExperiment: infrastructure for spatially-resolved transcriptomics data in R using Bioconductor. Bioinformatics 38, 3128–31 (2022).

[75] Beygelzimer, A., et al. FNN: fast nearest neighbor search algorithms and applications (2024). URL https://CRAN.R-project.org/package=FNN. R package version 1.1.4.1.

[76] Pedersen, T. L. ggraph: an implementation of grammar of graphics for graphs and networks (2024). URL https://CRAN.R-project.org/package=ggraph. R package version 2.2.1.

[77] Pedersen, T. L. tidygraph: a tidy API for graph manipulation (2024). URL https://CRAN.R-project.org/package=tidygraph. R package version 1.3.1.

78. CZI Cell Science Program et al. CZ CELLxGENE Discover: a single-cell data platform for scalable exploration, analysis and modeling of aggregated data. Nucleic Acids Res 53, D886–900 (2025).

