## Supplementary Material for "scDiagnostics: systematic assessment of cell type annotation in single-cell transcriptomics data"

**S3 Supplementary Discussion** **9**

**Supplementary Figures** **11**

**Supplementary Tables** **30**

### S1 Supplementary Methods

#### S1.1 Cell type annotation tools

Reference-based cell type annotation has become a standard approach in single-cell RNA-seq analysis, enabling researchers to transfer known cell type identities from well-annotated reference datasets to new query datasets. This strategy leverages the assumption that transcriptomically similar cells share the same biological identity. Multiple computational tools implement reference-based label transfer using different algorithmic approaches and distance metrics, each with distinct advantages and design philosophies. Here we focus on four widely-used annotation tools with mature implementations in R and Python: `SingleR` [1], `Azimuth` [2], `CellTypist` [3], and `scVI`/`scArches` [4, 5]. Many additional annotation tools exist in the literature [6]; the four tools presented here represent popular implementations that also provide confidence metrics or uncertainty estimates. Although most annotation tools provide per-cell confidence or uncertainty estimates reflecting internal algorithmic certainty, these metrics alone do not systematically validate dataset-level compatibility or diagnose the underlying causes of low-confidence assignments.

`SingleR` [1] performs annotation by computing Spearman correlations between each query cell and reference cell type pseudobulk profiles (aggregated expression per cell type), assigning each cell to the reference cell type with the highest correlation. A key feature is its built-in quality control mechanism: the "delta score" defined as the difference between the highest and second-highest correlation values provides a calibrated measure of annotation confidence. `SingleR` [1] also generates confidence score heatmaps visualizing correlations between cell types, enabling visual inspection of annotation ambiguity and potential mislabeling. Larger deltas indicate more unambiguous assignments, whereas small deltas suggest the cell is equidistant from multiple reference cell types. The tool can optionally flag and exclude cells with delta scores below a user-defined threshold, enabling systematic quality filtering.

`Azimuth` [2], implemented through `Seurat`'s label transfer framework [7], uses dimensionality reduction and weighted nearest-neighbor (wNN) graph-based methods [8]. It performs reference integration using anchor-based integration to align query cells to the reference principal component (PC) space, then transfers labels based on weighted k-nearest neighbor voting in the shared low-dimensional space. A prediction score is computed as the proportion of $k$-neighbors (typically $k = 30$) that vote for the predicted cell type, providing a confidence measure between 0 and 1.

`CellTypist` [3] uses a logistic regression-based classifier trained on reference data [9]. The method identifies genes consistently differentially expressed between cell types, then trains a multinomial logistic regression model. During prediction, confidence scores reflect the probability estimate from the logistic regression model for the predicted cell type. The tool supports ensemble prediction through majority voting across multiple similar models to improve prediction stability.

`scVI`/`scArches` [4,5] implements reference-based annotation using variational autoencoders (VAEs) [10]. The `scVI` framework trains a VAE on reference data to learn a low-dimensional latent representation that captures cell type variation. In this probabilistic framework, the model learns a distribution over possible latent representations for each cell rather than a single point estimate. The `scArches` [5] extension enables transfer learning [11] by adapting the reference model to query data through continued training on query cells while keeping most model parameters fixed. Uncertainty estimates derive from the learned posterior distribution, with cells mapping to intermediate regions between reference cell type clusters showing higher uncertainty.

#### S1.2 Limitations of existing cell type annotation tools

Although each of the four annotation tools provides useful per-cell confidence or prediction scores, relying on these metrics alone has several limitations. One important limitation is that they generally represent *relative* probabilities for the available reference cell types rather than *overall* structural fit between query and reference dataset. Annotation algorithms are optimized to determine which reference label is the best match compared to available alternative cell type labels. For instance, an inflamed monocyte clearly separated in high-dimensional space from healthy monocytes may still receive a near-optimal confidence score from a logistic regression classifier (e.g., `CellTypist` [3]) if it remains on the correct side of the decision boundary. Similarly, if an inflamed query monocyte is transcriptionally distant from a healthy reference monocyte, but even further from all other reference cell types (e.g., a T-cell), correlation-based methods (e.g., `SingleR` [1]) will still yield a large correlation delta and assign the monocyte label with high confidence. Consequently, standard confidence scores can confidently mask massive structural shifts driven by disease states, novel cell types, or technical batch effects.

`scDiagnostics` provides complementary diagnostics to address this issue by evaluating the *overall* structural fit of query cells to their assigned reference cell type. The package offers statistically sound, interpretable projection methods to assess how well query cells map to the reference, including PCA to capture overall variation patterns [12], discriminant analysis that highlights genes with the strongest cell type-differentiating power [13], and sliced inverse regression (SIR) to identify directions that optimally separate cell types and states [14]. The package also provides distributional alignment metrics using optimal transport and statistical distances (e.g., Bhattacharyya, Hellinger, Wasserstein [15–19]) to quantify the magnitude of structural divergence. To identify specific outlier cells, the package implements anomaly detection via isolation forests [20], coupled with downstream marker gene differential expression using random forest feature importance [21] to identify potential underlying drivers of detected anomalies.

Importantly, `scDiagnostics` is tool-agnostic: it requires only predicted cell type labels and log-normalized expression data in standard `SingleCellExperiment` format [22], making it compatible with annotations from any method. By applying `scDiagnostics` as a post-hoc auditing layer, users can jointly interpret standard relative confidence scores alongside overall anomaly scores. This allows analysts to distinguish true high-quality matches from biologically shifted or technically distorted cells, gaining mechanistic insight into annotation quality without replacing their established annotation workflows.

As a complementary baseline to this density-based approach, the package also provides an anomaly diagnostic based on principal component analysis (PCA) reconstruction error. For a given cell type, this method computes a local PCA space using only the reference cells, projects the query cells into it, and attempts to reconstruct their original gene expression profiles. Anomalies are then flagged by calculating the sum of squared errors (SSE) between the original and reconstructed profiles. Because the reference PCA only models biological variation present in the reference dataset, a high reconstruction error identifies query cells that exhibit novel gene expression patterns (such as a disease state) that the reference model cannot explain.

### S2 Supplementary Results

#### S2.1 Quantitative benchmarking and parameter sensitivity

To systematically benchmark anomaly detection with `scDiagnostics` across common challenges encountered in real single-cell data, we utilized the Zeisel brain scRNA-seq dataset [23]. We split the data into reference and query subsets and systematically withheld specific reference populations to construct three distinct out-of-distribution (OOD) scenarios (Figure S3A): a distinct glial lineage (Astrocytes), one of two closely related pyramidal neuronal cell subtypes, and a rare cell type (Microglia). Using `SingleR` [1], query cells from these missing populations were forced into incorrect reference categories. To rigorously evaluate the application of Isolation Forests (IF) in detecting these forced misannotations, we implemented a simple and intuitive baseline method based on PCA Reconstruction Error (PCAR). Interestingly, our evaluations revealed that this baseline displays certain complementary aspects to IF: while IF models structural density within the principal signal space, PCAR measures deviations in the null space (i.e., calculating the sum of squared reconstruction residuals from a localized PCA manifold).

At baseline, both the IF diagnostic and the PCAR baseline accurately identified the misannotated OOD cells (AUROC $> 0.85$, Figure S3B).

**Performance across confounder gradients.** To simulate suboptimal mapping conditions, we subjected the query datasets to controlled gradients of annotation label noise (0–40% shuffled labels), progressive class imbalance (reducing reference populations from up to 300 cells down to 10 cells), and simulated batch effects (mean shifts ranging from 0 to +2 log-normalized counts). Anomaly detection with `scDiagnostics` retained strong discriminatory power across moderate noise and batch levels (Figure S3C–H). In addition to AUROC, Sensitivity, and Specificity, we explicitly evaluated the Area Under the Precision-Recall Curve (AUPRC), which provides a more robust performance metric for highly imbalanced scenarios, such as the detection of the rare Microglia population (Figure S4). Specificity remained above 0.8 in most scenarios, empirically demonstrating that the algorithms control false positive rates and effectively ignore in-distribution sparse regions (Figure S6).

**Comparison of IF to the PCAR baseline.** Comparing IF performance against the PCAR baseline showed that these two approaches are to a certain extent complementary. Because its partitioning algorithm effectively captures global structural density within the principal signal space, IF excelled at detecting distinct and rare populations. This was particularly evident for the rare Microglia population, where IF maintained near-perfect AUPRC across noise and batch gradients, vastly outperforming the PCAR baseline (Figure S4F–H). Conversely, PCAR provided a more consistent baseline sensitivity for detecting the highly related Pyramidal SS neurons under severe noise and imbalance (Figure S5C–D; Figure S4C–E). Because PCAR measures strict geometric deviations from localized reference manifold rules (null space) rather than global density, it serves as a highly effective complementary baseline for resolving very subtle shifts between closely related cellular subtypes.

**Feature space selection.** To ensure compact representation of high-dimensional single-cell data prior to anomaly detection, we evaluated the impact of different feature spaces (Global PCA vs. Targeted HVGs) using parameter grid searches (Figure S7). By projecting cells into a dense, high-signal subspace, `scDiagnostics` effectively neutralizes the "curse of dimensionality" before anomaly detection occurs. The appropriate choice of this subspace depends on the dataset's variance structure. When anomalies are subtle or driven by few genes (such as the closely related pyramidal neuron states; Figure S3A), running IF on a restricted set of Targeted HVGs yields higher specificity (Figure S7, blue markers). Conversely, when anomalies are driven by broad transcriptional shifts (e.g., the COVID-19 and MERFISH disease states), Global PCA effectively captures the dominant variance. To accommodate diverse topologies, `scDiagnostics` supports anomaly detection on either representation.

**Adaptive anomaly thresholds.** Our parameter grid searches demonstrated that adapting anomaly thresholds to the reference population provides a more consistent balance of sensitivity and specificity across varying data topologies than arbitrary fixed cutoffs (e.g., an absolute threshold of 0.5; see Figures S7 and S8). Under this adaptive approach, `scDiagnostics` first computes baseline anomaly scores for the reference cells. The algorithm then calculates the median and the median absolute deviation (MAD) of these reference scores, establishing a data-adaptive threshold defined as the reference median plus a scalar multiple of the MAD. As established by widely adopted single-cell workflows such as the *Orchestrating Single-Cell Analysis* (OSCA) framework [22], utilizing MAD-based adaptive thresholds is a common best practice for robustly handling dataset-specific variance. Because baseline anomaly scores naturally fluctuate depending on the intrinsic variance of a given cell type or the dimensionality of the dataset, fixed thresholds often fail to generalize across datasets. By automatically calibrating the boundary to the specific reference distribution, this MAD-based approach alleviates the need for manual threshold tuning. As shown in our grid searches, applying a two- or three-MAD multiplier consistently optimized detection performance across varying cell types and dataset complexities (Figures S7 and S8).

#### S2.2 Application of scDiagnostics to the output of major annotation tools (scRNA-seq COVID-19 PBMC atlas)

To assess the complementarity of scDiagnostics with established annotation workflows, we applied four widely used tools—SingleR [1], Azimuth [2], CellTypist [3], and scArches [5]—to the COVID-19 peripheral blood mononuclear cell (PBMC) dataset from Stephenson et al. [24]. Consistent with the main text, our analysis focused on CD14$^+$ monocytes to evaluate the detection of known disease-driven transcriptomic dysregulation.

**Overview of annotation outputs and confidence distributions.** All four annotation methods assigned CD14$^+$ monocyte identity (Figure S9A, C, E, G), with consistent spatial clustering in UMAP space despite methodological differences. However, the methods showed substantial variation in confidence score distributions (Figure S9B, D, F, H). SingleR [1] and scArches [5] produced lower mean confidence scores compared to Azimuth [2], while CellTypist [3] exhibited near-uniform high confidence score across cells. This diversity in confidence metrics underscores the importance of orthogonal validation approaches.

**Interferon gene dysregulation in query cells.** Consistent with severe COVID-19 pathobiology, CD14$^+$ monocytes displayed marked elevation of interferon genes (IFN signature: 25 genes), visible as distinct high-scoring clusters in UMAP space (Figure S10). This IFN activation was heterogeneous across cell clusters, indicating variable innate immune responses even within the CD14$^+$ monocyte compartment.

**PCA structure and anomaly detection.** We projected query cells into the principal component space of a healthy reference and applied scDiagnostics anomaly detection (Figure S11). For each annotation method, the PCA landscape revealed structured separation between query and reference CD14$^+$ monocytes, consistent with transcriptomic remodeling in COVID-19 disease. Anomalous cells identified by scDiagnostics were clustered in PCA space and often co-embedded with cells displaying elevated IFN signatures. This correspondence suggests that scDiagnostics captures genuine transcriptomic divergence rather than noise.

**Relationship between annotation confidence and anomaly status.** We tested whether annotation tool confidence inversely correlates with anomaly designation. Distribution analysis (Figure S12) revealed that anomalous cells consistently displayed lower annotation confidence compared to non-anomalous cells across all four methods (Table S3). Wilcoxon rank-sum tests showed highly significant differences ($p < 0.001$ for all methods). Notably, SingleR [1] exhibited significant distribution shifts despite identical medians, indicating that anomalous cells occupy a distinct confidence distribution even when central tendencies match. This highlights the sensitivity of scDiagnostics to nuanced changes in the confidence score distribution. Crucially, we observed that a substantial proportion of anomalous cells still received high confidence scores from the underlying annotation tools (Figure S12). As detailed conceptually in Section S1.2, this discrepancy arises from the fundamental difference between relative classification probabilities and overall structural fit. Because annotation tools are optimized to find the best relative match among available categories, they can confidently assign a label to a diseased cell as long as it remains closer to its target lineage than to any alternative. By explicitly measuring absolute structural deviation, scDiagnostics successfully flags these biologically shifted cells that standard tools confidently mask.

**IFN gene elevation in anomalous and low-confidence cells.** To identify cells at the extremes of annotation confidence while retaining statistical power, we stratified each method's cells based on its score direction: for methods where higher scores indicate greater confidence (SingleR [1], Azimuth [2], CellTypist [3]), high-confidence cells were defined as those in the $\geq$ 75th percentile and low-confidence cells as those in the $\leq$ 25th percentile. For scArches [5], which uses uncertainty scores (lower is better), the interpretation reverses: high-confidence cells are those in the $\leq$ 25th percentile (lowest uncertainty) and low-confidence cells are those in the $\geq$ 75th percentile (highest uncertainty). We then calculated mean IFN signature scores for each stratified group (Figure S11). Both low-confidence and anomalous cell populations showed marked elevation of the IFN signature relative to reference cells (Table S2), with median IFN scores in anomalous cells 1.5–2-fold higher than non-anomalous cells across all methods (Wilcoxon, $p < 0.001$). Importantly, low-confidence cells also displayed elevated IFN signatures, suggesting that annotation tool uncertainty reflects underlying biological dysregulation rather than a technical artifact.

**Concordance of gene expression shifts across annotation tools.** To assess whether annotation confidence and `scDiagnostics` anomaly detection identify overlapping transcriptomic dysregulation, we computed $\log_2$-fold changes of 25 IFN genes in low-confidence versus high-confidence cells, and in anomalous versus non-anomalous cells (Figure S13). Spearman rank correlation of these gene shift vectors revealed high concordance across methods (Table S4): `Azimuth` [2]($r = -0.910$), `scArches` [5] ($r = -0.920$), and `CellTypist` [3] ($r = -0.642$), all $p < 0.001$. The negative correlations reflect the relationship between annotation confidence scores and `scDiagnostics` anomaly scores: annotation confidence scores increase with certainty, while `scDiagnostics` anomaly scores increase with divergence. `SingleR` [1] showed weaker correlation ($r = 0.182$, $p = 0.38$), likely due to its compressed, near-binary confidence distribution. Overall, these correlations indicate substantial overlap in transcriptomic features driving both annotation confidence and anomaly detection.

**Interpretation and implications.** These analyses demonstrate that `scDiagnostics` provides orthogonal and complementary insights to annotation tool confidence scores. Rather than simply assigning or rejecting one of the available reference cell types to a query cell, `scDiagnostics` identifies transcriptomic inconsistencies between query and reference populations. Notably, `scDiagnostics` anomalies do not perfectly align with low annotation confidence, suggesting the method detects reference-query mismatches that annotation tools' confidence scores fail to capture. Annotation confidence reflects internal algorithmic certainty about cell identity, while `scDiagnostics` provides explicit additional evidence (via PCA structure, distributional shifts, and coordinated gene expression changes) for transcriptomic divergence. The consistency of anomaly patterns across four different annotation tools indicates that detected anomalies reflect genuine biology rather than method-specific artifacts. Both annotation confidence and anomaly status strongly associate with elevated IFN signatures, supporting the biological relevance of both signals in identifying dysregulated monocytes in severe COVID-19 infection.

#### S2.3 Application of `scDiagnostics` to the output of major annotation tools (MERFISH timecourse of DSS-induced colitis)

To demonstrate the compatibility of `scDiagnostics` with major annotation tools also for a spatially-resolved single-cell dataset, we applied the four annotation tools— `SingleR` [1], `Azimuth` [2], `CellTypist` [3], and `scArches` [5]— to the mouse colitis timecourse dataset from Cadinu et al. [25] (Figure S14). Annotations were transferred from healthy (day 0) reference to DSS-treated (day 9) query samples. In the corresponding section of the main manuscript, inflamed cell states were collapsed into quiescent categories prior to `scDiagnostics` application. Here, we retained inflamed annotations and applied `scDiagnostics` to each tool's outputs, using author-annotated inflamed fibroblasts as ground truth.

**Annotation tool discordance in inflammation-associated fibroblasts (IAFs).** All four annotation methods assigned cell type identity with largely concordant clustering in UMAP space (Figure S15A, C, E, G). However, annotation tools showed substantial disagreement when classifying cells labeled as IAFs in the ground truth. `SingleR` [1], `CellTypist` [3], and `scArches` [5] assigned most IAFs to the Fibroblast category. `Azimuth` [2] and `scArches` [5] assigned larger proportions of Smooth Muscle Cells (SMCs): 532 of 923 cells for `Azimuth` [2] (57.6%), and 313 of 1024 cells for `scArches` [5] (30.6%) (Figure S16). This pattern was consistent across samples and indicates systematic misclassification of a subset of IAFs.

**PCA structure and ECM gene expression.** Fibroblasts (both reference and query) were projected into principal component space using all ECM homeostasis genes (*Col1a2*, *Timp2*, *Col6a1*, *Sparc*,

*Dpt*) (Figure S17A–C). Query cells were clearly separated from reference cells across the first three PCs for all four tools, reflecting transcriptomic differences between healthy and inflamed states. Query fibroblasts displayed reduced ECM homeostasis scores compared to reference cells, with mean reductions visible in the color gradients. This pattern was observed regardless of which annotation method assigned the cells.

**Anomaly detection results.** We applied `scDiagnostics` anomaly detection using isolation forests [20] to fibroblasts annotated by each method (Figure S17B–D). Cells annotated as SMC by `Azimuth` [2]showed high anomaly rates: 532 of 540 cells (98.5%) were detected as anomalous. `scArches` [5] showed 313 of 338 SMC-annotated cells (92.6%) as anomalous. Cells annotated as Fibroblast showed lower anomaly rates: `Azimuth` [2]16 of 24 (66.7%), `scArches` [5] 1 of 2 (50.0%).

**ECM gene dysregulation by annotation category.** We computed $\log_2$-fold changes of the five ECM genes stratified by annotation assignment within each method (Figure S18). All query fibroblasts showed significant ECM suppression relative to reference ($p < 0.001$ across methods). Cells annotated as SMCs showed the most pronounced suppression, with mean $\log_2$-fold changes ranging from $-0.8$ to $-1.2$ for *Col1a2* and *Col6a1*. Cells annotated as Fibroblast retained higher ECM expression but still showed suppression relative to reference.

**Marker gene expression patterns.** We identified the top 10 SMC-specific and fibroblast-specific marker genes from the healthy reference (FDR $< 0.01$) and visualized their expression across four cell groups (Figure S19). Cells annotated as SMC expressed intermediate levels of SMC markers (*Igfbp5*, *Grem1*, *Flnc*) and reduced fibroblast markers (*Col1a2*, *Col18a1*, *Mmp2*). Reference SMCs and fibroblasts showed expected marker separation with minimal overlap. This intermediate expression pattern in query cells suggests partial overlap with SMC gene signature rather than true smooth muscle identity.

***Col1a2* expression and anomaly status.** *Col1a2* expression differed significantly between anomalous and non-anomalous cells across all annotation methods (Table S5). Anomalous cells showed reduced mean expression (`SingleR`: $0.233 \pm 0.664$ vs. $1.977 \pm 1.184$; `Azimuth`: $0.212 \pm 0.672$ vs. $2.109 \pm 1.196$), with mean differences of $-1.744$ to $-1.897$ units (all $p < 0.001$, Student's $t$-test). Non-anomalous cells retained higher *Col1a2* levels despite being drawn from inflamed tissue.

**Consistency across methods.** The patterns of annotation discordance, high anomaly rates in SMC-assigned cells, and ECM suppression were consistent across all four annotation methods. This reproducibility across independent methods suggests that the transcriptomic pattern is not a method-specific artifact but reflects genuine biological differences between cell groups.

**Summary.** `scDiagnostics` identified systematic differences between cells assigned to different categories by the four annotation methods. Cells mislabeled as SMC displayed high anomaly rates (92.6%–98.5%) and marked ECM gene suppression. These cells show a combination of reduced fibroblast marker expression and intermediate SMC marker expression, indicating a dysregulated intermediate state rather than clear SMC identity. The strong association between anomaly detection and reduced *Col1a2* expression confirms that `scDiagnostics` captures transcriptomic dysregulation in inflamed fibroblasts. The consistency of these patterns across four independent annotation methods supports the validity of the detected anomalies.

#### S2.4 Runtime and memory usage

To evaluate the computational runtime and memory usage under different scenarios, we first benchmarked the main diagnostic functions of `scDiagnostics` on the COVID-19 PBMC dataset from Stephenson et al. [24], comprising 23,201 healthy reference cells and 48,148 COVID-19 query cells after quality control (18,472 genes). Benchmarks were performed using the `bench` R package [26] on a laptop equipped with a 13th Gen Intel® Core™ i7-13800H processor (2.50 GHz) and 32 GB RAM. On this dataset, the global PCA step on the reference (`processPCA`, 1,000 HVGs) completed in 5.4 seconds and used 0.27 GB of RAM. Projecting the full query dataset into the reference PCA space (`projectPCA`, first 5 PCs) required

2.2 seconds and 2.7 GB of RAM. Isolation-forest-based anomaly detection (`detectAnomaly`; 1,000 HVGs, 500 trees, MAD thresholding) completed in 1.3 seconds with a peak memory allocation of 0.86 GB, while the PCA reconstruction-error diagnostic (`calculateReconstructionError`; 100 local HVGs, first 5 PCs, MAD thresholding) completed in 0.82 seconds using 0.64 GB. The most computationally intensive function, `calculateGeneShifts`, which combines anomaly detection with pseudo-bulk differential expression and automated loading-vector selection, completed in 10.7 seconds and used 3.2 GB of RAM for CD14$^+$ monocytes. These results indicate that, for a typical case–control analysis with tens of thousands of cells, all diagnostics can be run interactively on a standard laptop.

**Scaling to atlas-level datasets.** To assess behavior on larger datasets, we additionally benchmarked `scDiagnostics` on HDF5-backed COVID-19 PBMC data loaded directly from the CELLxGENE portal. We randomly sampled increasing total numbers of cells (10,000, 50,000, 100,000, 250,000, and 500,000 cells), splitting each sample into 30% reference and 70% query cells, and focused on the most abundant reference cell type at each scale. For each sample, we measured execution time and peak RAM usage for `processPCA`, `projectPCA`, `detectAnomaly`, `calculateReconstructionError`, and `calculateGeneShifts` (Figure S2). Across this range, all routines exhibited approximately linear growth in runtime and memory as a function of the total number of cells. Even at 500,000 total cells, the most expensive steps (`processPCA`, `projectPCA`, and `calculateGeneShifts`) completed within tens of seconds, and peak RAM usage remained below 21 GB. Of note, anomaly detection was particularly lightweight: both `detectAnomaly` and `calculateReconstructionError` completed in under 5 seconds and used less than 2 GB of memory at the largest scale.

**Practical recommendations.** These results suggest several straightforward strategies for applied users analyzing large-scale datasets. First, because the primary computational cost lies in the initial dimensionality reduction, we recommend pre-computing the global reference PCA using a fixed subset of highly variable genes (e.g., 1,000 HVGs). Supplying this pre-computed object directly to the downstream diagnostic functions amortizes the cost and significantly reduces overall runtime. Second, for very large datasets ($>$ 1M cells) with highly imbalanced cell type distributions, users can apply random subsampling for highly abundant cell types prior to fitting the reference manifold. This approach caps the number of cells used for model training, which is typically sufficient to stabilize diagnostic scores while drastically reducing peak memory requirements. Finally, because `scDiagnostics` seamlessly integrates with Bioconductor's out-of-core data representations, we highly recommend passing large datasets as HDF5-backed `SingleCellExperiment` [22], `SpatialExperiment` [27], or `SpatialFeatureExperiment` [28] objects via the `DelayedArray` framework [29] to maintain a minimal memory footprint. By following these recommendations, users can maintain excellent performance on standard laptops even when evaluating hundreds of thousands of cells.

### S3 Supplementary Discussion

**Comparison to existing tools.** To compare `scDiagnostics` to existing computational tools we considered four categories of tools based on their primary functionality (Table S1): standard reference mapping (e.g., `SingleR` [1]), out-of-distribution (OOD) detection (e.g., `scArches` [5]), disease-state discovery (e.g., `Augur` [30]), and post-hoc diagnostics (e.g., `scDiagnostics`). While standard reference mapping tools provide relative confidence scores, and specialized deep-learning models offer OOD detection based on complex neural network architectures, `scDiagnostics` fills a unique niche. It is designed to be algorithm-agnostic, allowing users to retain their preferred established annotation workflows while providing an independent, statistical assessment of absolute structural divergence and automated identification of the underlying gene shifts.

**Consistency across annotation tools.** In both the COVID-19 and MERFISH datasets, anomaly detection and downstream gene expression comparisons yielded overlapping results regardless of the underlying annotation tool (`SingleR` [1], `Azimuth` [2], `CellTypist` [3], or `scArches` [5]). This consistency across tools suggests that the detected structural deviations are primarily driven by features of the expression data rather than artifacts introduced by a specific annotation algorithm.

**Relationship to confidence scores.** Annotation confidence scores reflect an algorithm's internal certainty regarding a label assignment. In the COVID-19 analysis, while anomalous cells frequently exhibited lower annotation confidence, the overlap was not complete. This indicates that structural anomaly scores measure a property distinct from classification confidence. Utilizing both metrics may therefore provide a more comprehensive view of annotation behavior (Figure S1A), particularly when confidence score distributions vary widely across different tools (e.g., the near-uniform high confidence of `CellTypist` versus the broader distribution of `SingleR`).

**Interpreting anomalies.** Detecting a statistical anomaly does not automatically define its cause. However, coupling anomaly detection with gene expression analysis can assist in interpretation (Figure S1B). In the MERFISH dataset, stratifying cells by anomaly status highlighted differences in ECM gene expression and SMC marker levels. While this provides descriptive context for why certain fibroblasts deviated from the reference, identifying the underlying biological or technical mechanisms requires further domain-specific investigation.

**Limitations.** The application of `scDiagnostics` is constrained by several factors. First, it relies on the availability of an annotated reference; its outputs are directly dependent on the quality and composition of that reference [31]. Second, as noted above, structural anomalies do not inherently distinguish between novel cell types or states, disease-induced transcriptional shifts, or technical batch effects. Differentiating these requires careful review of the experimental design and context-dependent considerations. Third, as the field starts moving toward large-scale foundation models [32–34], paradigms for cell type annotation will likely evolve, though (accordingly adapted) post-hoc diagnostic frameworks will continue to serve an important role in evaluating model generalization to unseen datasets [35, 36].

**Practical guidelines.** When applying `scDiagnostics`, investigators may benefit from using multiple annotation tools to assess whether anomalies are tool-dependent or generalize across tools. Upon identifying anomalous populations, reviewing marker gene expression, PCA projections, and available metadata (such as disease condition, spatial location, or sample batch) is necessary to determine whether the deviation likely reflects a technical artifact or a biological signal. We recommend the following decision criteria for jointly interpreting `scDiagnostics` anomaly scores alongside standard annotation confidence metrics (Supplementary Figure S1):

- *High anomaly scores accompanied by marker alignment failures and strong correlations with batch or QC metrics:* Suspect a technical artifact (e.g., unmodeled batch effects, high ambient RNA, or poor-quality cells).

- *High anomaly scores concentrated in a latent region lacking reference neighbors:* Suspect reference incompleteness (i.e., a true out-of-reference population or novel cell state absent from the atlas).

- *Low annotation confidence paired with low anomaly scores:* Suspect annotation or model failure. The query cell resides within the reference manifold but sits on an ambiguous boundary between two cell types (likely a safe annotation; review marker genes to confirm).

- *High annotation confidence paired with high anomaly scores and a coherent pathway signature:* Suspect genuine biological drift. The cell belongs to the assigned lineage but is undergoing a disease-driven state change (e.g., IFN activation in monocytes or ECM suppression in fibroblasts).

#### Supplementary Figures

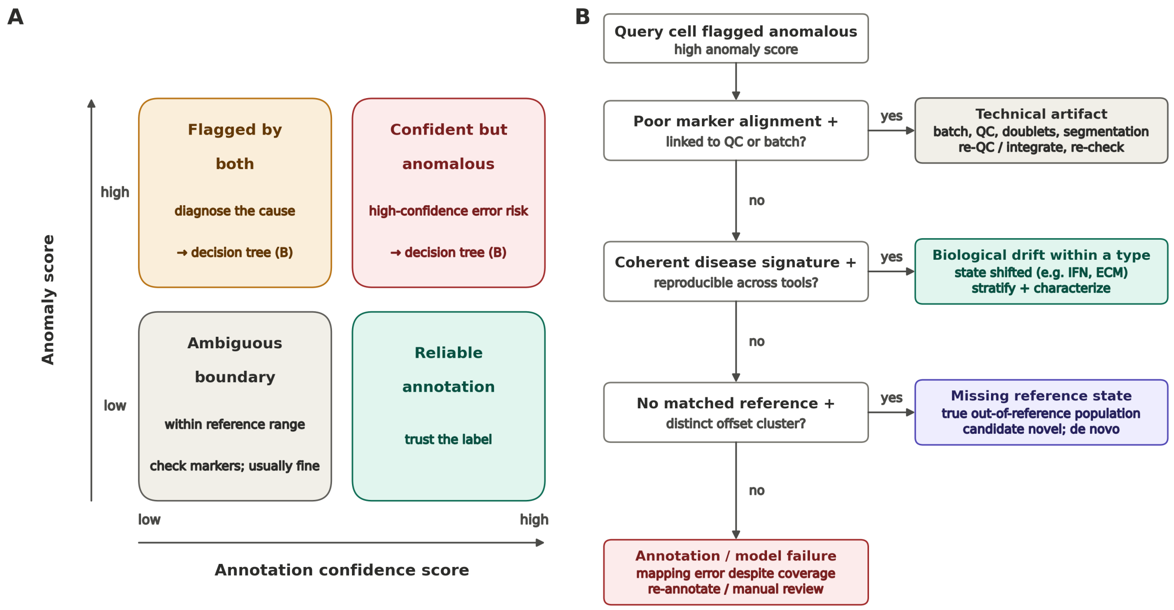

**Figure S1:** Interpretation of cell type annotation anomalies detected with `scDiagnostics`. **(A)** Joint interpretation of the per-cell anomaly score computed by `scDiagnostics` and the confidence (or prediction) score reported by the annotation tool of choice. Cells with low anomaly scores (bottom) are consistent with the reference: high-confidence assignments are reliable (bottom right), whereas low-confidence assignments (bottom left) typically reflect ambiguity at the boundary between related reference cell types rather than an out-of-reference population, and can be resolved by inspecting marker genes. High anomaly scores (top) warrant further diagnosis regardless of confidence; cells assigned with high confidence yet flagged as anomalous (top right) represents the high-risk case in which standard workflows silently misassign transcriptomically divergent cells. **(B)** Decision framework for identifying potential causes for a high-anomaly query population using complementary diagnostics implemented in the package. Technical artifacts are primary confounders and present as poor marker-gene alignment that correlates with quality-control metrics or batch structure, and are addressed by re-inspection or integration. A coherent, interpretable expression signature that reproduces across annotation tools indicates biological drift within an existing cell type—a shifted state to be analyzed and characterized separately rather than discarded. Anomalous cells forming a distinct cluster with no matched reference counterpart indicate a genuine out-of-reference (missing) population and a candidate for *de novo* annotation. When none of these apply despite adequate reference coverage, the anomaly is likely a result of annotation or model failure, warranting re-annotation or manual review.

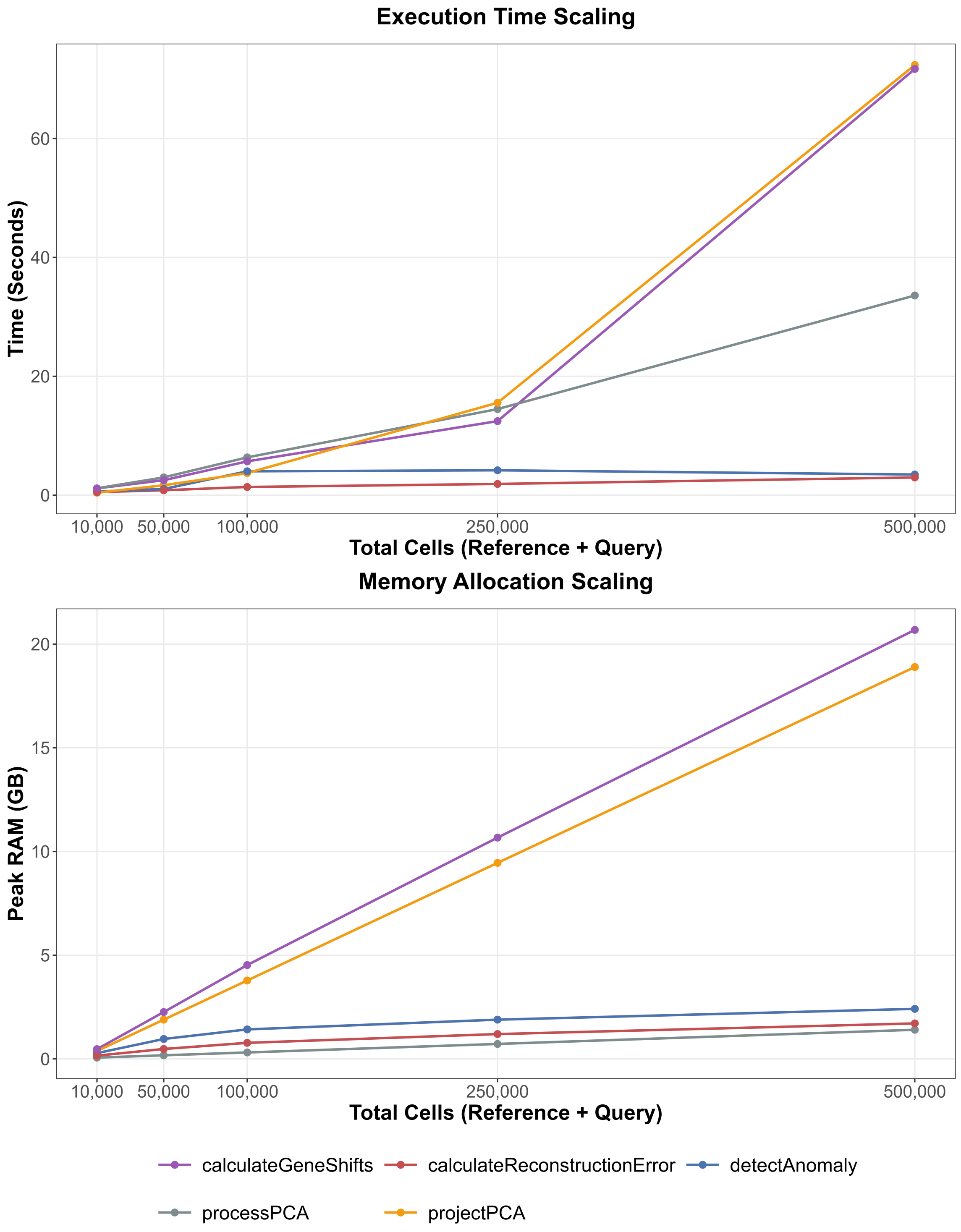

**Figure S2:** Runtime and memory usage of the main functions of `scDiagnostics` on COVID-19 PBMC data [24]. Left: Execution time ($y$-axis) for main diagnostic functions on an increasing total number of combined reference and query cells ($x$-axis), using subsamples of 10,000, 50,000, 100,000, 250,000, and 500,000 cells drawn from the COVID-19 PBMC dataset. Right: Corresponding peak RAM usage for each function. Colors denote individual `scDiagnostics` functions: PCA on the reference (`processPCA`, grey), projection of query cells onto the reference PCA space (`projectPCA`, orange), isolation-forest anomaly detection (`detectAnomaly`, blue), PCA reconstruction-error diagnostics (`calculateReconstructionError`, red), and combined anomaly + differential expression analysis (`calculateGeneShifts`, purple). The evaluation was carried out on a standard laptop (13th Gen Intel Core i7, 32 GB RAM).

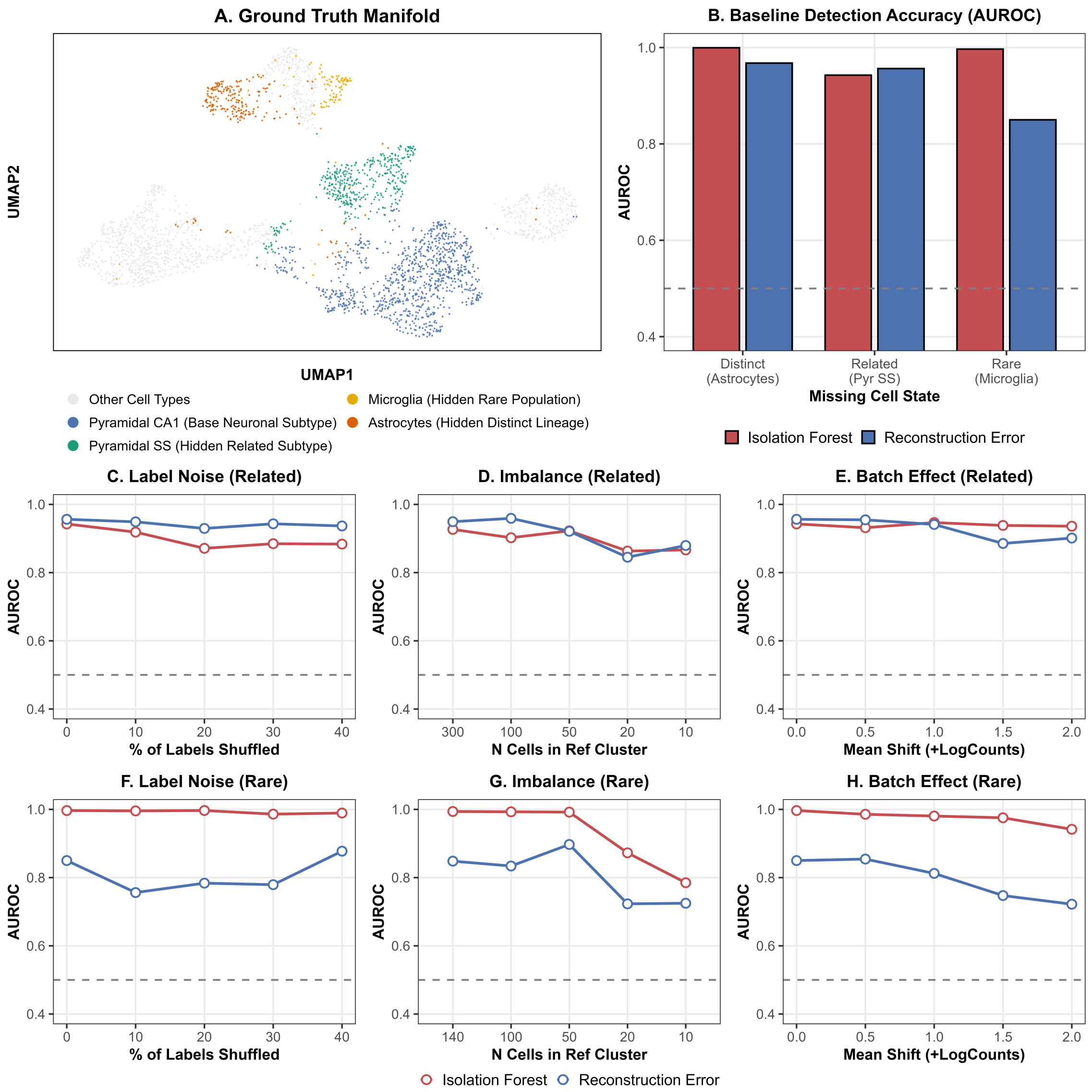

**Figure S3:** AUROC benchmarking of anomaly detection under realistic single-cell confounders. (A) UMAP visualization of the Zeisel brain dataset [23] highlighting the evaluated reference (Pyramidal CA1) and three withheld query populations: a distinct glial lineage (Astrocytes), a closely related neuronal subtype (Pyramidal SS), and a rare population (Microglia). (B) Baseline AUROC for the Isolation Forest diagnostic (red) compared against the PCA Reconstruction Error baseline (blue) when detecting the hidden populations. (C–H) AUROC evaluated across gradients of annotation label noise (0–40% shuffled labels), progressive class imbalance (reducing the reference cluster from 300 down to 10 cells), and simulated batch effects (mean log-count shifts from 0 to 2) for both the closely related subtype (C–E) and the rare population (F–H).

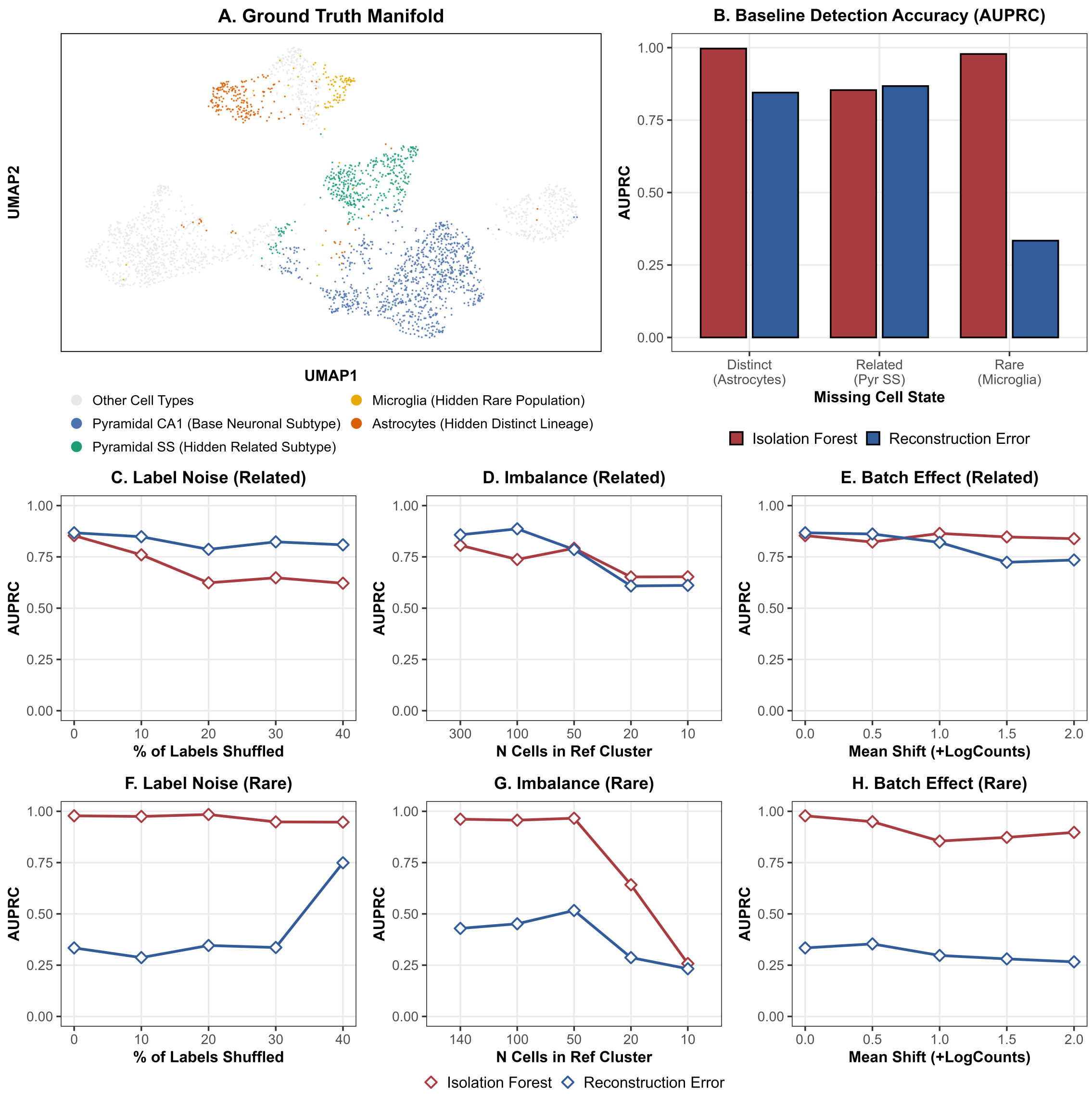

**Figure S4:** AUPRC (Area Under the Precision-Recall Curve) benchmarking of anomaly detection under realistic single-cell confounders. Corresponding to the scenarios in Figure S3, panels evaluate the AUPRC of the Isolation Forest diagnostic (red diamonds) and the PCA Reconstruction Error baseline (blue diamonds). AUPRC provides a highly robust performance metric for evaluating detection accuracy in severely imbalanced scenarios, such as the detection of the rare Microglia population. (A) UMAP visualization of the evaluated populations. (B) Baseline AUPRC for detecting the hidden query populations. (C–H) AUPRC evaluated across gradients of annotation label noise (0–40%), progressive class imbalance (300 down to 10 cells), and simulated batch effects (0 to 2 log-count mean shift) for both the closely related subtype (C–E) and the rare population (F–H).

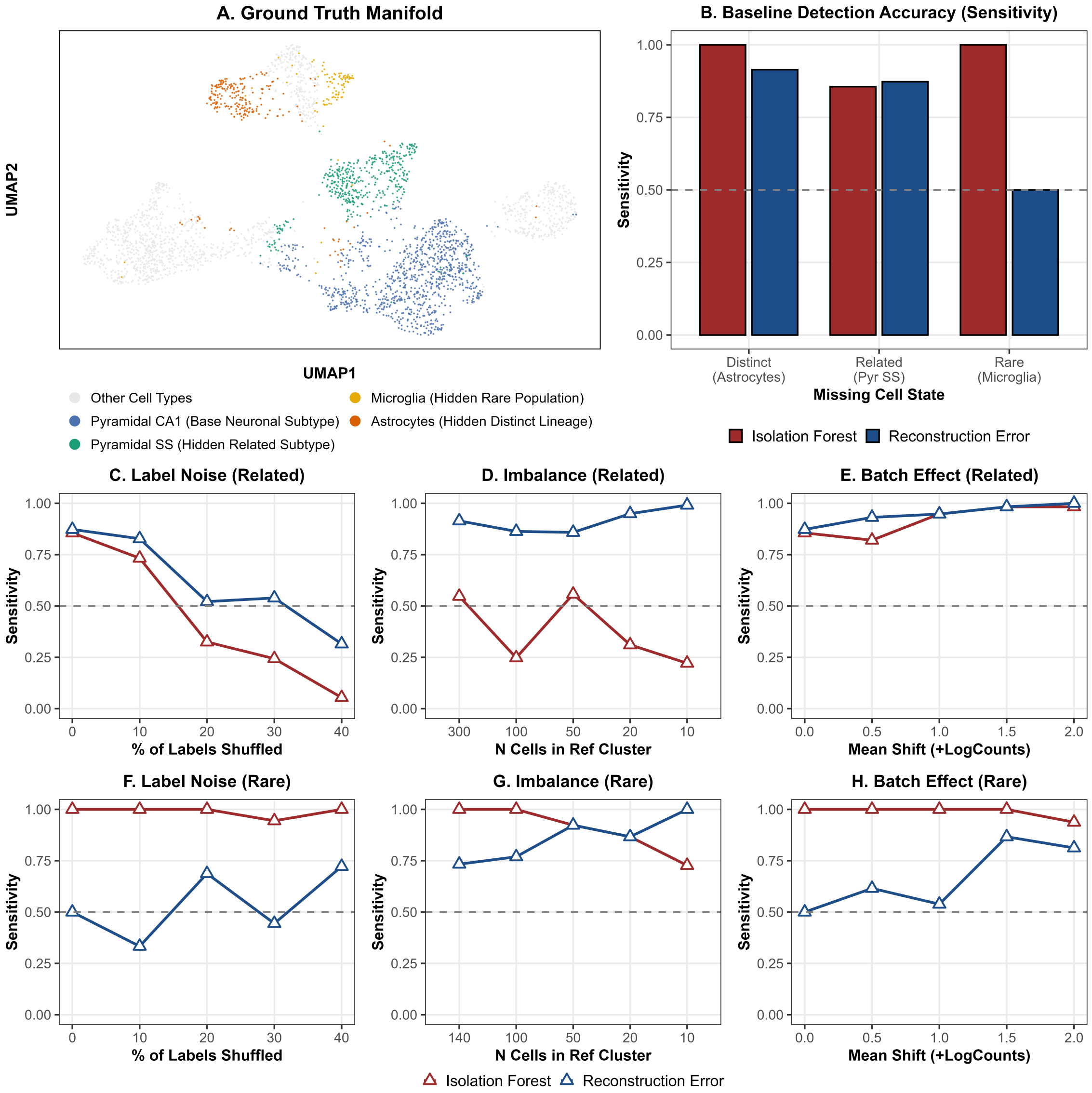

**Figure S5:** Sensitivity (True Positive Rate) benchmarking of anomaly detection. Corresponding to the scenarios in Figure S3, panels evaluate the Sensitivity of the Isolation Forest diagnostic (red triangles) and the PCA Reconstruction Error baseline (blue triangles) in correctly identifying true out-of-distribution (OOD) query cells across gradients of annotation label noise (C, F), class imbalance (D, G), and simulated batch effects (E, H).

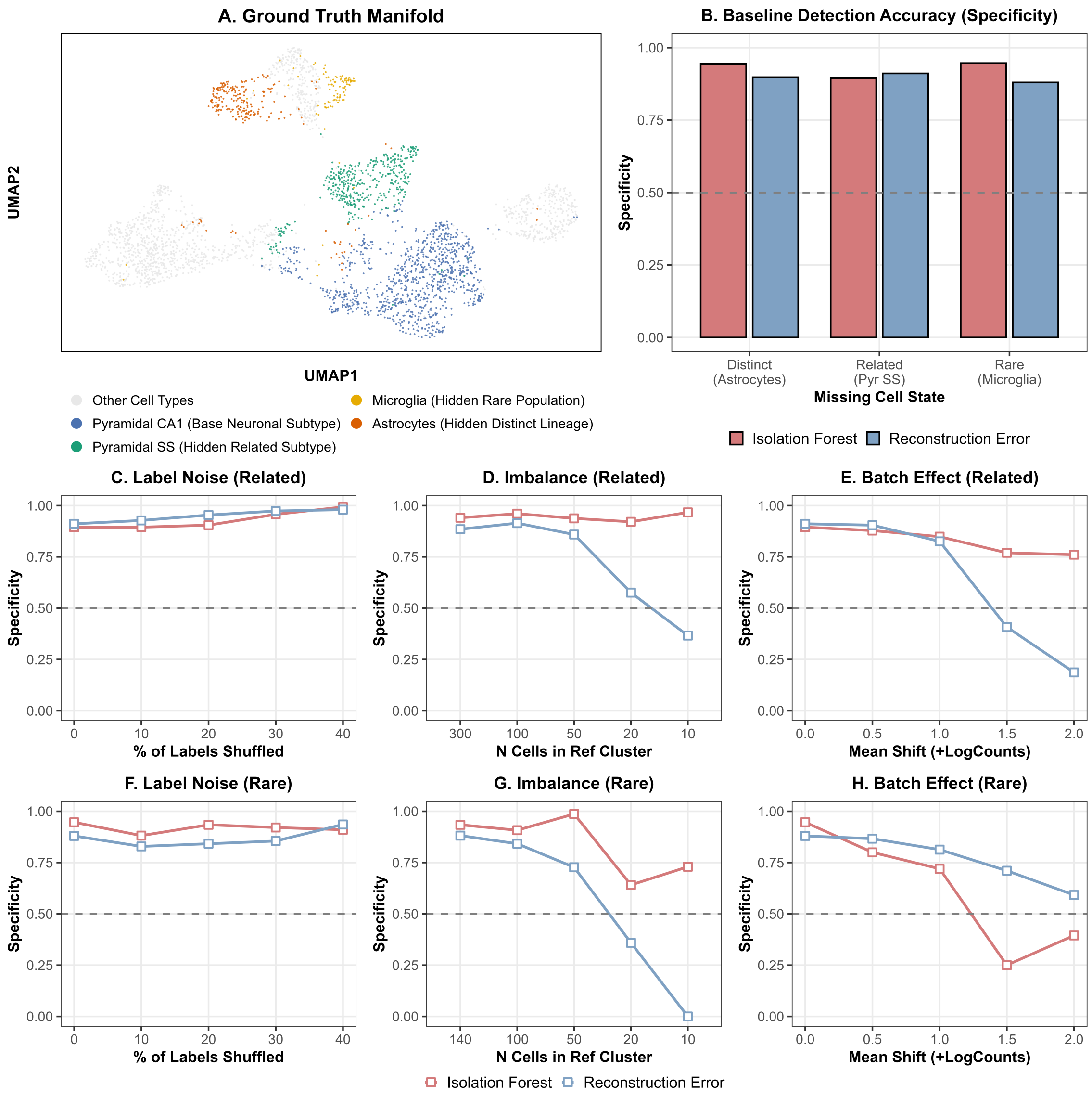

**Figure S6:** Specificity (True Negative Rate) benchmarking of anomaly detection. Corresponding to the scenarios in Figure S3, panels evaluate the Specificity of the Isolation Forest diagnostic (red squares) and the PCA Reconstruction Error baseline (blue squares) in correctly ignoring in-distribution, reference-like query cells across gradients of annotation label noise (C, F), class imbalance (D, G), and simulated batch effects (E, H). Sustained high specificity demonstrates that the algorithms effectively control the false positive rate and are not biased by sparse high-dimensional regions.

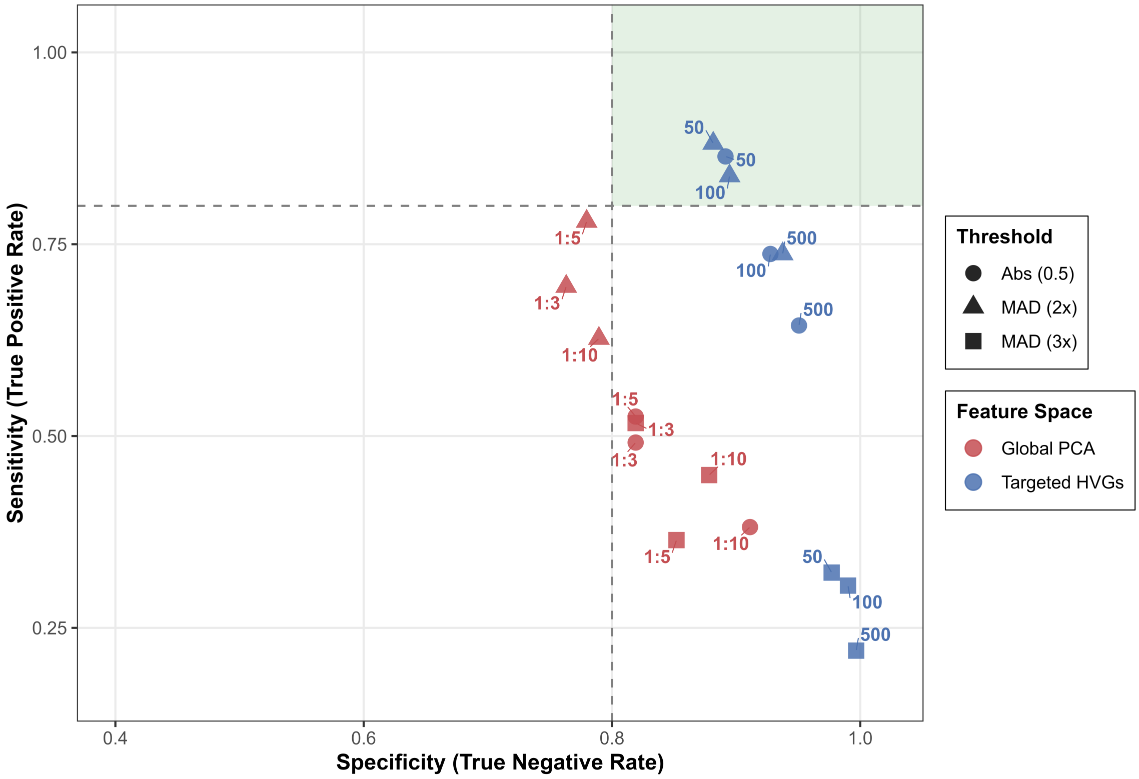

**Figure S7:** Parameter sensitivity and feature space tuning for the Isolation Forest diagnostic. Grid search evaluating Isolation Forest performance when detecting the hidden closely related neuronal subtype (Pyramidal SS). Models were run using either a Global PCA subspace (red) or a restricted set of Targeted HVGs (blue). To demonstrate the stability of data-adaptive thresholds, the algorithm was evaluated across an arbitrary fixed threshold of 0.5 (circles) versus dynamic cutoffs based on the Median Absolute Deviation (MAD) of the reference baseline (2x MAD, triangles; 3x MAD, squares). The green shaded region highlights the optimal performance zone where both Sensitivity and Specificity are $> 0.8$.

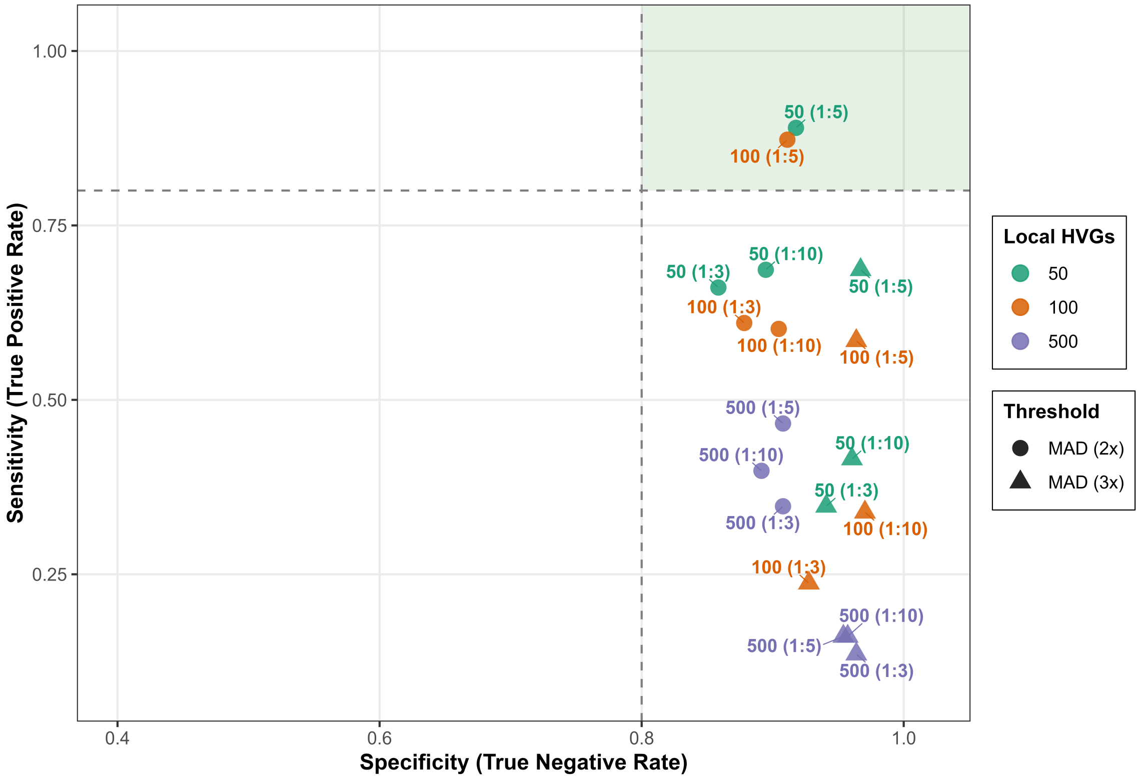

**Figure S8:** Parameter sensitivity tuning for the PCA Reconstruction Error baseline. Grid search evaluating the PCA Reconstruction Error diagnostic when detecting the hidden closely related neuronal subtype (Pyramidal SS). The algorithm was tested across combinations of highly variable genes (HVGs: 50, 100, 500) and the number of principal components (PCs: 1-3, 1-5, 1-10), utilizing data-adaptive threshold multipliers (2x MAD, 3x MAD). The green shaded region highlights the optimal performance zone where both Sensitivity and Specificity are $> 0.8$.

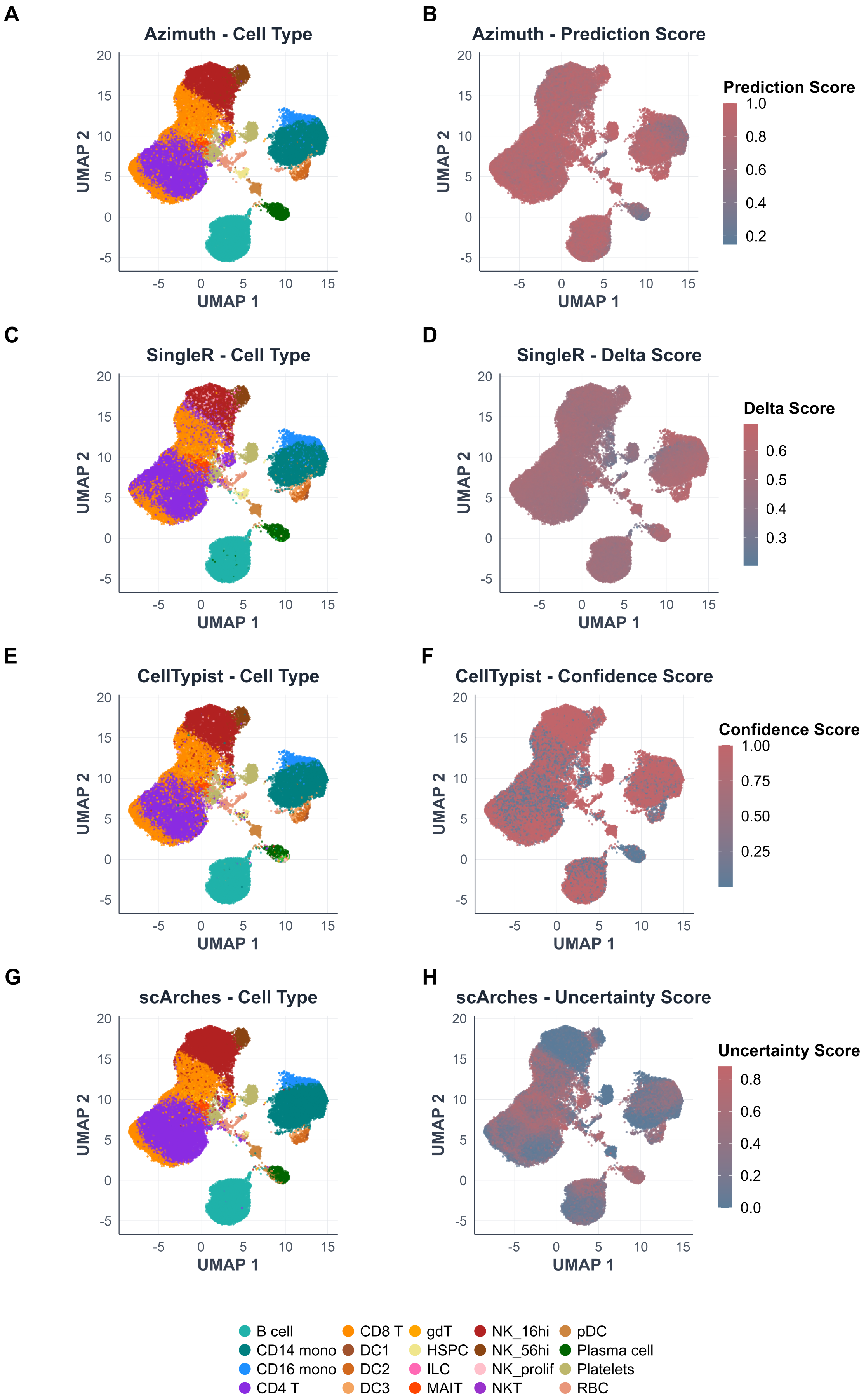

**Figure S9:** Cell type annotations and confidence scores across four annotation methods in COVID-19 $CD14^{+}$ monocytes. UMAP plots of PBMCs colored by (A, C, E, G) cell type predictions from `Azimuth` [2], `SingleR` [1], `CellTypist` [3], and `scArches` [5], or (B, D, F, H) method-specific confidence scores (prediction score for `Azimuth` [2], Delta score for `SingleR` [1], confidence score for `CellTypist` [3], and uncertainty score for `scArches` [5]). Left panels display distinct cell type clusters including myeloid, lymphoid, and erythroid lineages. Right panels show confidence score distributions, with blue indicating lower confidence (or higher uncertainty for `scArches` [5]) and red indicating higher confidence.

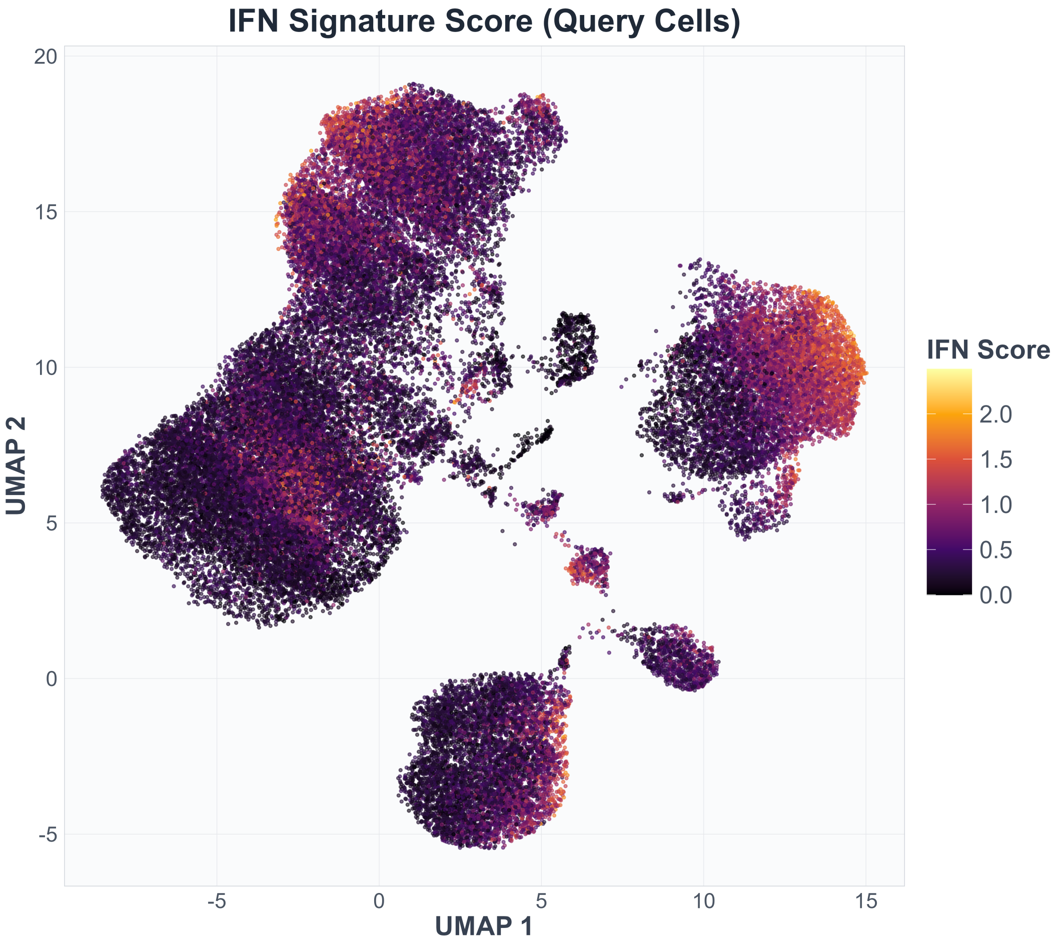

**Figure S10:** Interferon gene signature expression across COVID-19 PBMC. UMAP visualization showing mean expression of 25 interferon genes (IFN signature from Yoshida et al. [37]). Cells are colored by IFN signature score (mean $\log_2$-normalized expression), with yellow/orange indicating high IFN expression and black indicating low expression.

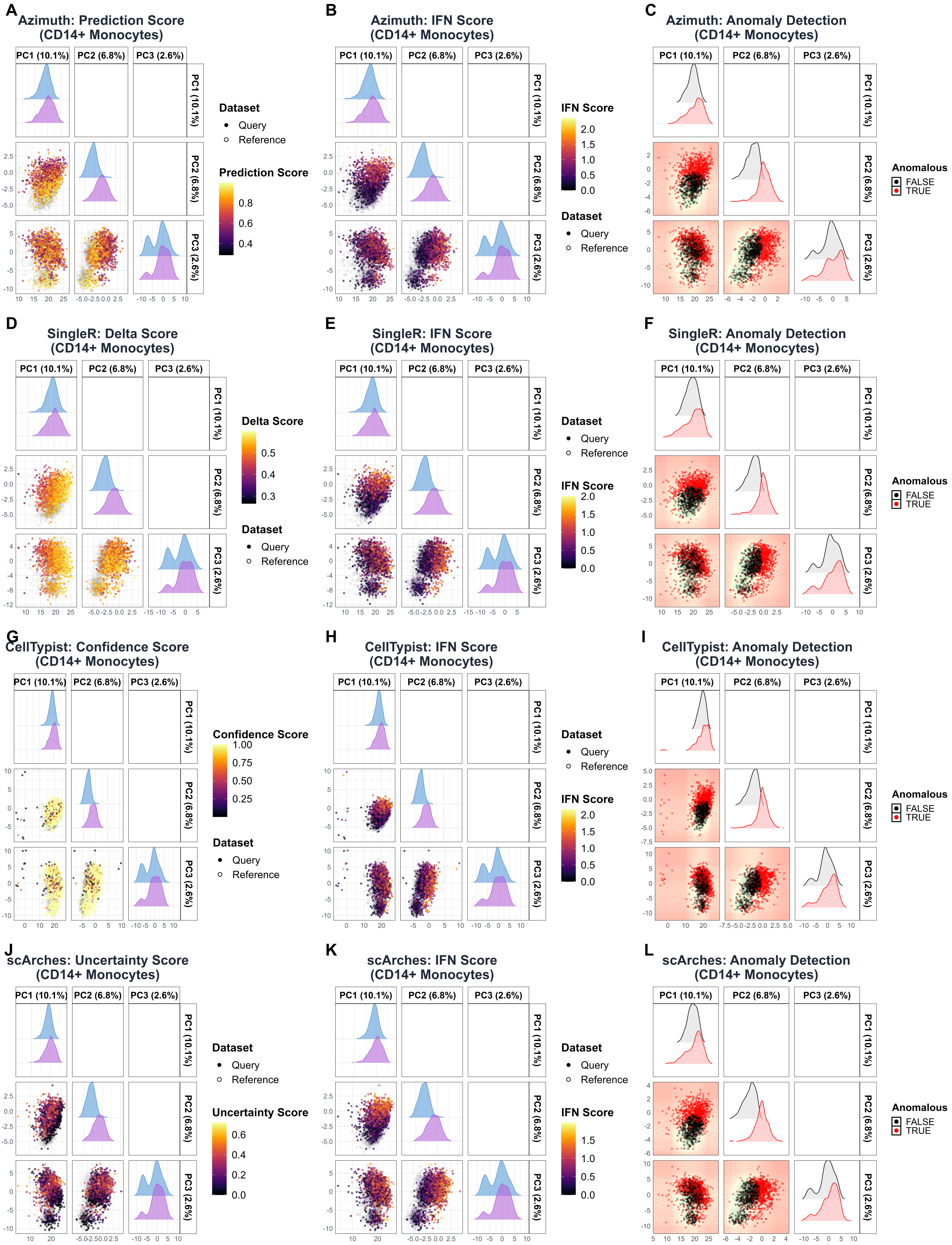

**Figure S11:** Principal component analysis (PCA) and anomaly detection for CD14$^+$ monocytes across annotation methods. (A–C) `Azimuth` [2]: PCs colored by prediction score (left), IFN signature score (middle), and `scDiagnostics` anomaly classification (right). (D–F) `SingleR` [1], (G–I) `CellTypist` [3], and (J–L) `scArches` [5] shown similarly, with `SingleR` [1], `CellTypist` [3], and `scArches` [5] using Delta score, confidence score, and uncertainty score, respectively. Reference cells (circles) and query cells (filled points) are plotted in PCA space. Anomalous cells identified by `scDiagnostics` are highlighted in red in the right panels for each method.

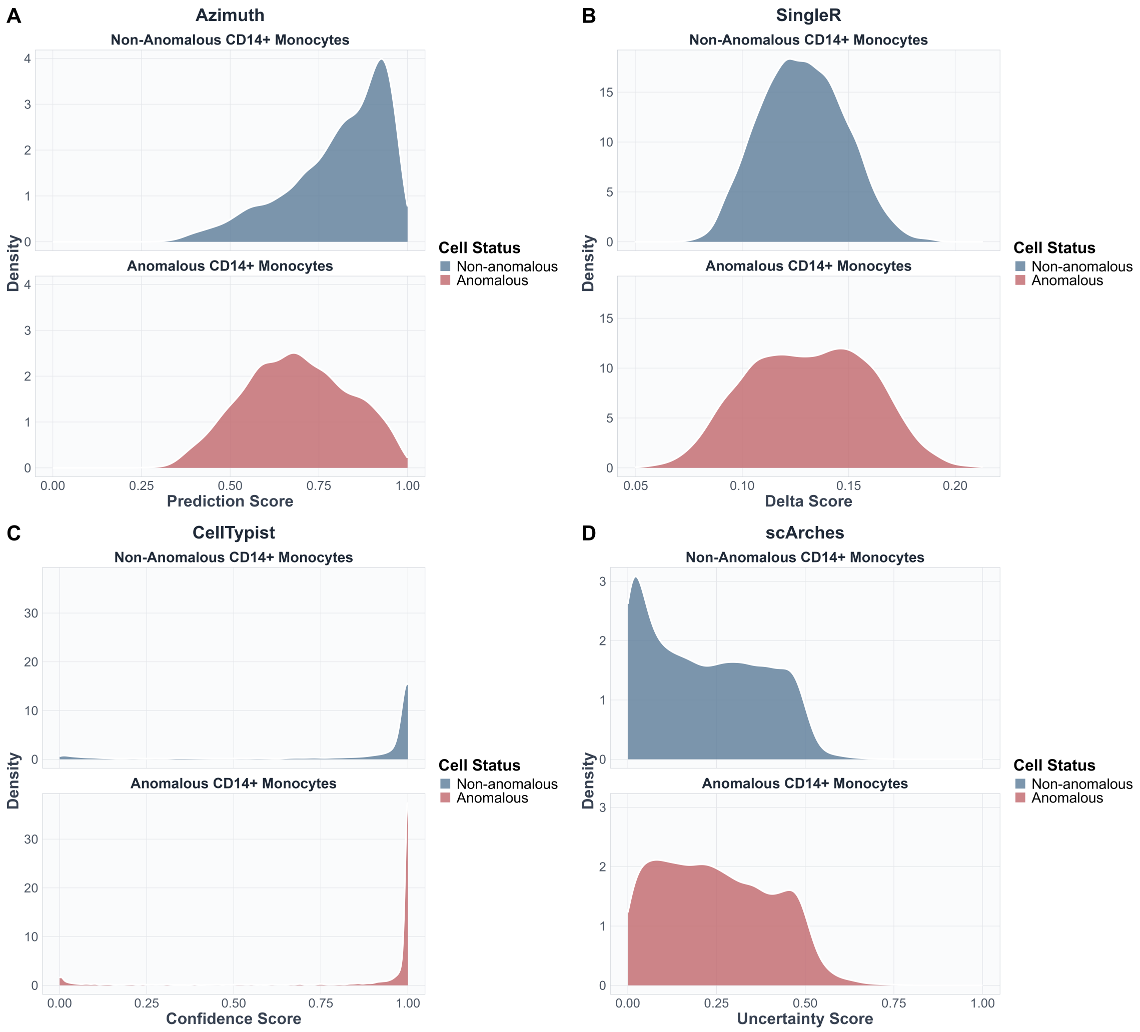

**Figure S12:** Distribution of annotation tool confidence scores stratified by anomaly status in $CD14^+$ monocytes. Density plots comparing score distributions between anomalous (top) and non-anomalous (bottom) $CD14^+$ monocytes for (A) `Azimuth` [2] prediction scores, (B) `SingleR` [1] Delta scores, (C) `CellTypist` [3] confidence scores, and (D) `scArches` [5] uncertainty scores. Distributions are separated by cell status (non-anomalous in blue, anomalous in red), with higher density indicating more cells at a given score.

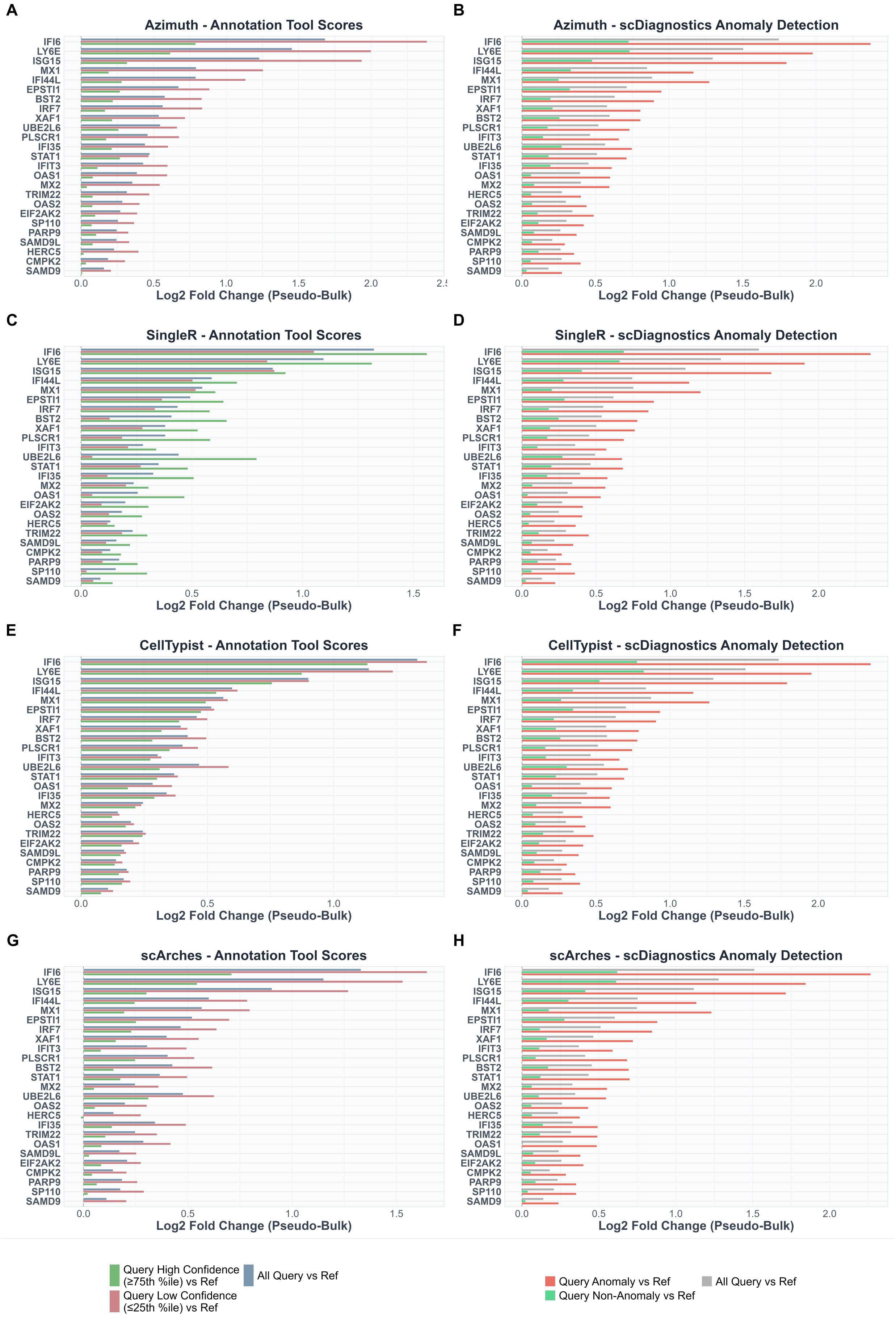

**Figure S13:** Gene expression shifts in interferon signature genes across annotation methods and stratification approaches. For each annotation method (rows: `Azimuth`, `SingleR`, `CellTypist`, `scArches`), $\log_2$-fold changes of 25 interferon genes are shown for cells stratified by annotation tool confidence quantiles (left column) and by `scDiagnostics` anomaly classification (right column). Left plots compare gene expression in high-confidence (green), low-confidence (red), and all query cells (gray) relative to reference. Right plots compare anomalous (red), non-anomalous (green), and all query cells (gray) relative to reference. Genes are ordered by statistical significance ($p_{\text{adj}} < 0.05$).

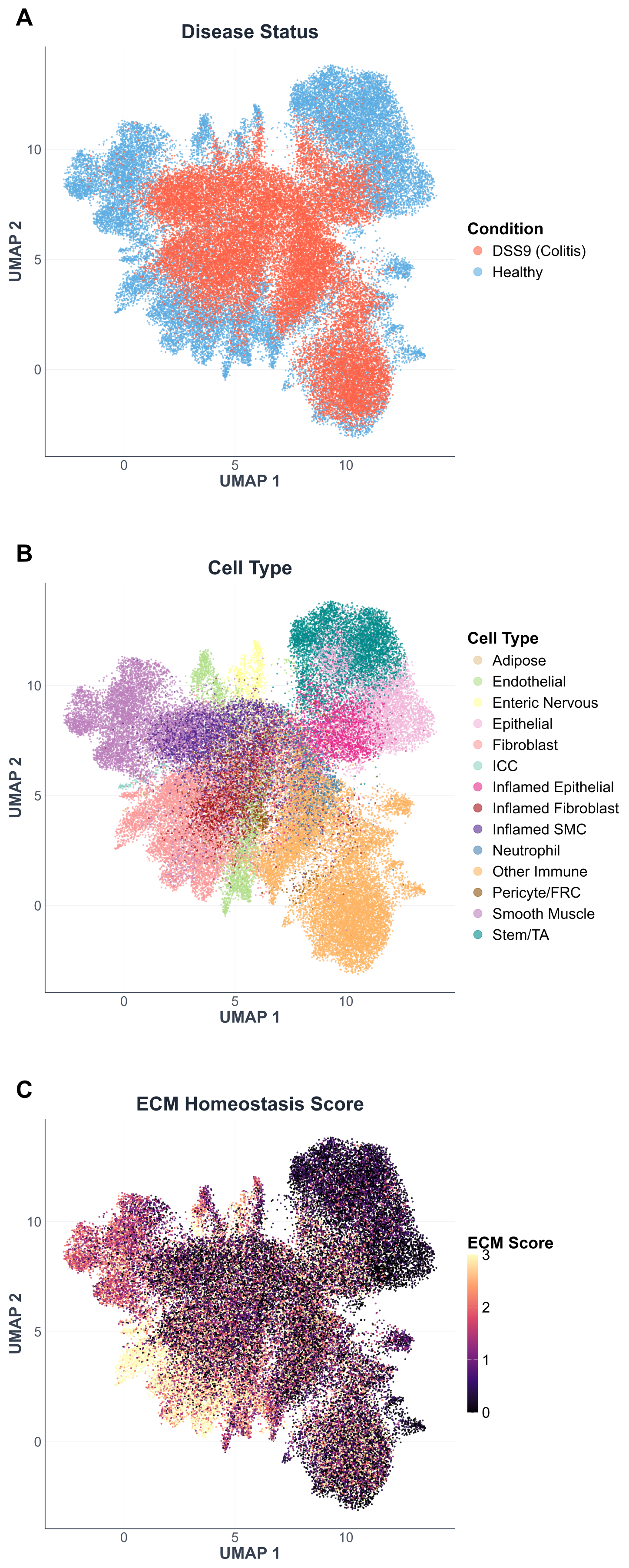

**Figure S14:** UMAP visualization of healthy and DSS9-induced colitis colon samples showing (A) disease status, (B) cell type annotations, and (C) ECM homeostasis score computed from mean expression of *Col1a2*, *Timp2*, *Col6a1*, *Sparc*, and *Dpt*.

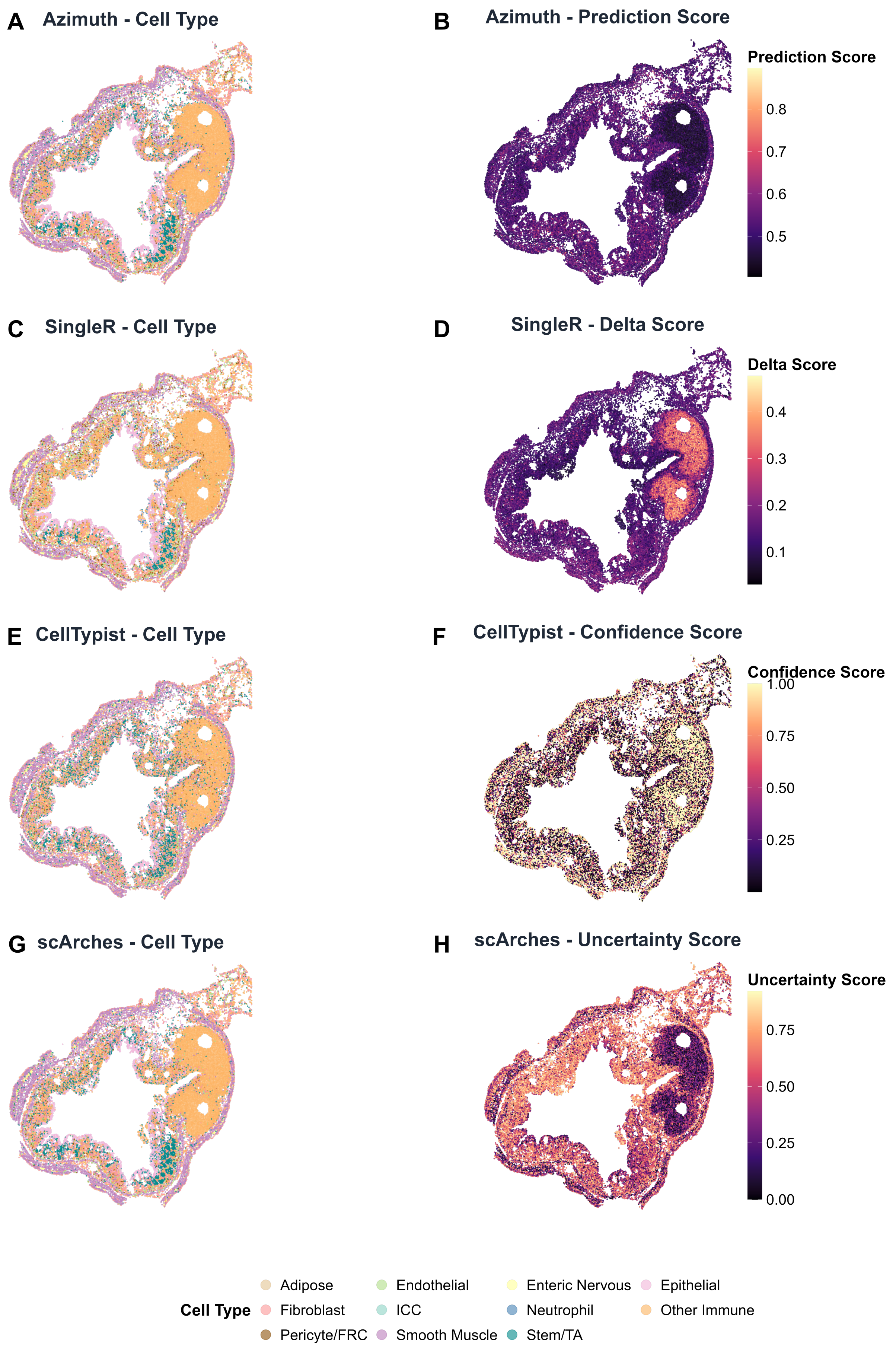

**Figure S15:** Spatial profiling of mouse colon tissue slices annotated with four cell type annotation tools: `Azimuth` [2] (A-B), `SingleR` [1] (C-D), `CellTypist` [3] (E-F), and `scArches` [5] (G-H). Left column shows predicted cell type assignments; right column shows corresponding confidence/prediction scores.

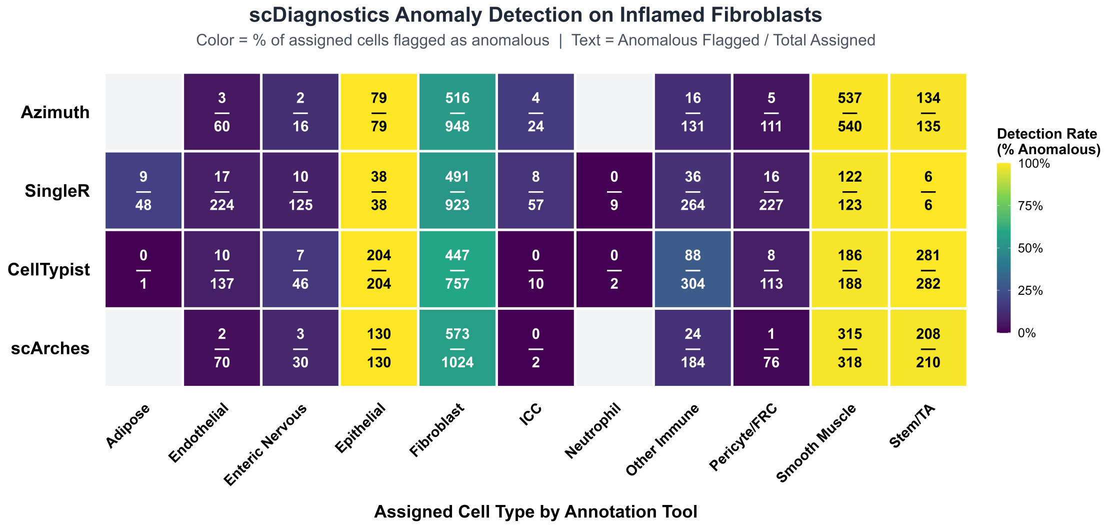

**Figure S16:** Heatmap showing the distribution of ground truth IAFs across cell type assignments predicted by four annotation methods (`Azimuth` [2], `SingleR` [1], `CellTypist` [3], and `scArches` [5]). Cell color intensity represents the percentage of assigned cells successfully flagged as anomalous by `scDiagnostics` (detection rate). Categories with zero assigned cells are shaded in light gray to reduce visual clutter. Text within each cell indicates the number of anomalous cells detected by `scDiagnostics` (numerator) divided by the total number of IAFs assigned to that cell type (denominator).

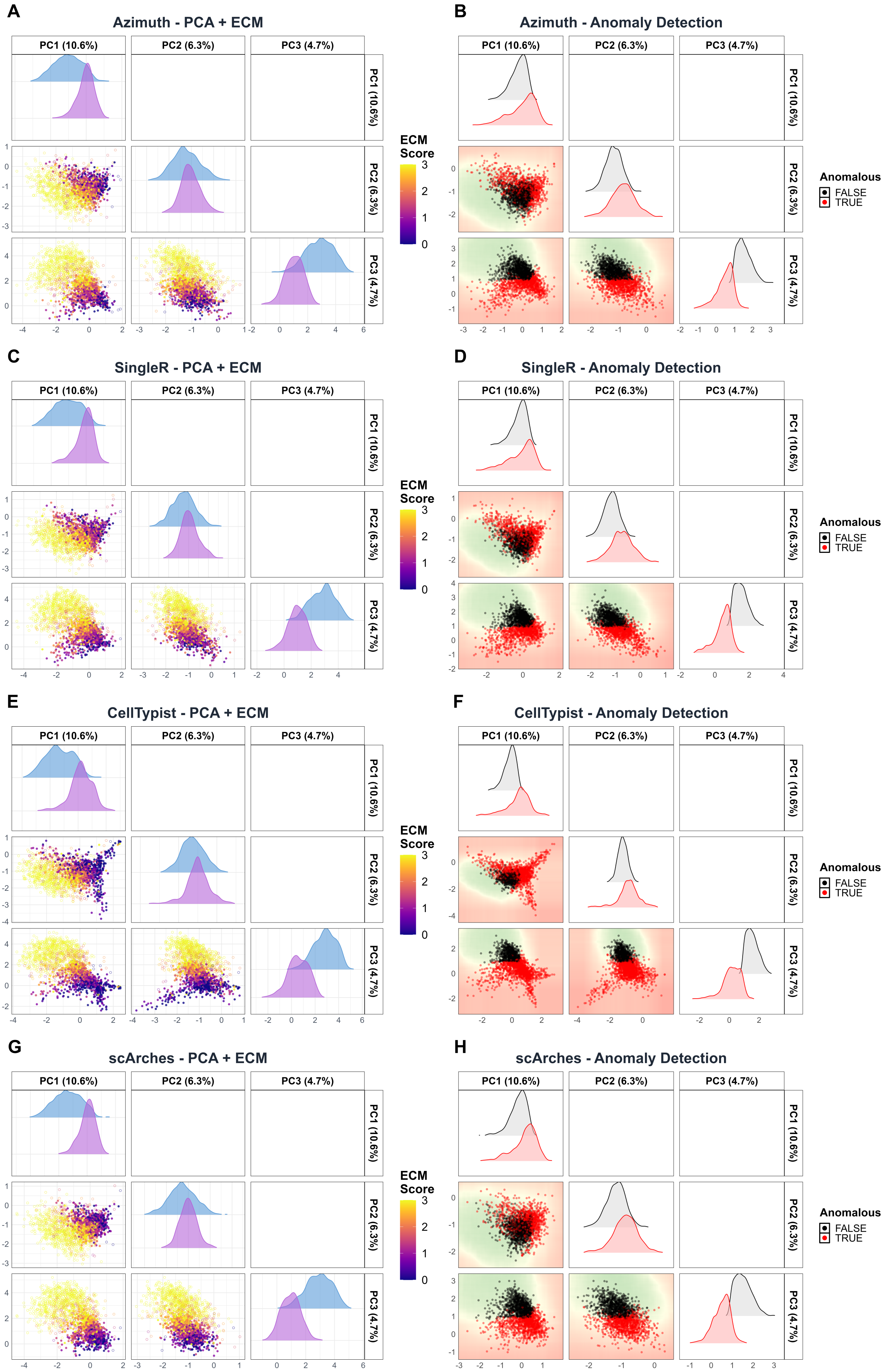

**Figure S17:** Fibroblast analysis across four annotation methods (`Azimuth` [2], `SingleR` [1], `CellTypist` [3], `scArches` [5]) applied to DSS9 samples. Left column shows PCA projections of healthy reference (open circles) and DSS9 query (filled circles) fibroblasts, colored by ECM homeostasis score. Right column shows anomaly detection results using isolation forests, with anomalous cells highlighted in red.

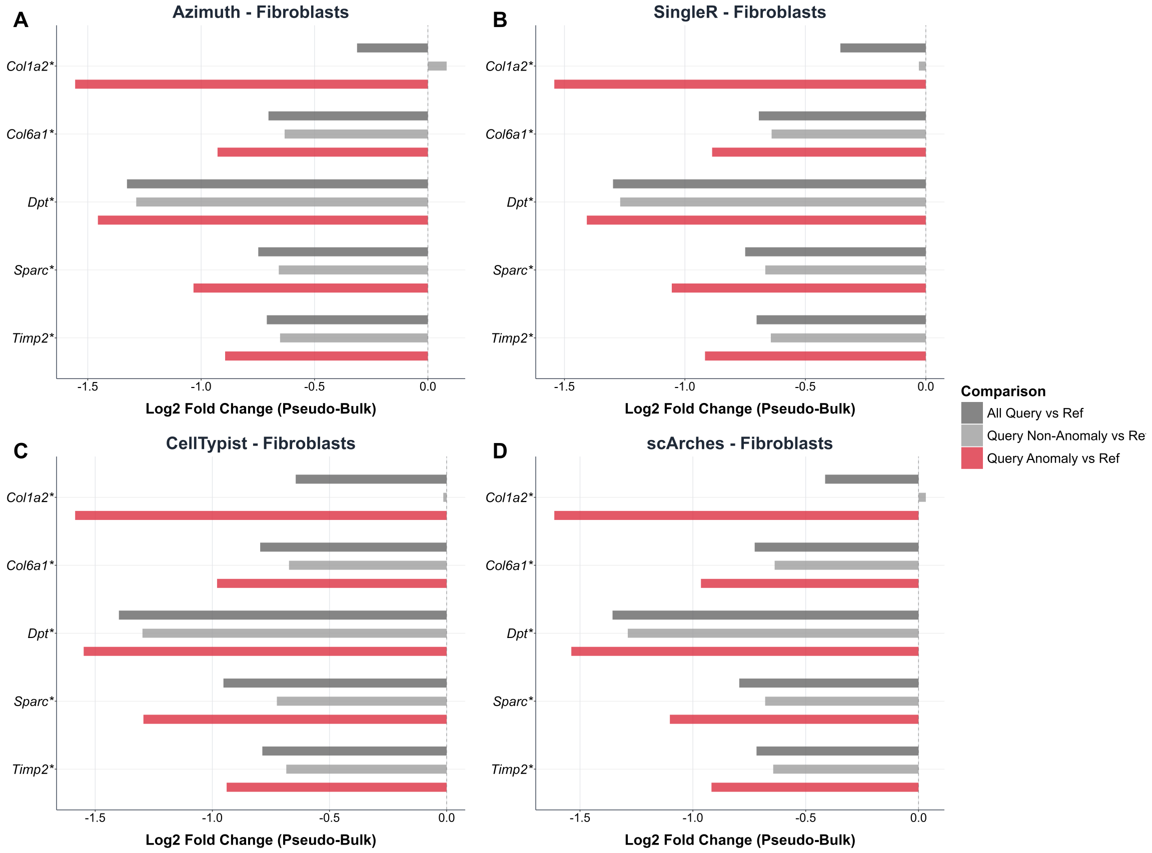

**Figure S18:** $\text{Log}_2$-fold changes of five ECM homeostasis genes (*Col1a2*, *Timp2*, *Col6a1*, *Sparc*, *Dpt*) in DSS9 fibroblasts compared to healthy reference. Bars show comparisons across all query cells versus reference (gray), non-anomalous query cells versus reference (light red), and anomalous query cells versus reference (dark red).

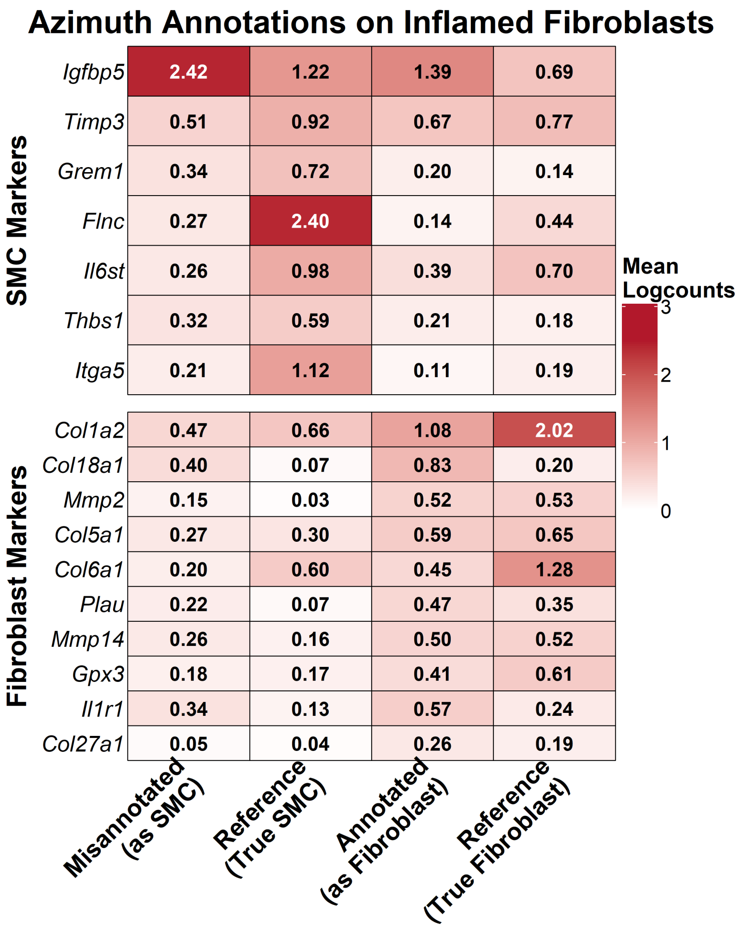

**Figure S19:** Heatmap showing mean log-normalized expression of the top 10 SMC and top 10 fibroblast marker genes across four cell groups: inflamed fibroblasts misannotated as smooth muscle cells (Misannotated as SMC), healthy reference SMCs (Reference True SMC), inflamed fibroblasts correctly annotated as fibroblasts (Annotated as Fibroblast), and healthy reference fibroblasts (Reference True Fibroblast).

### Supplementary Tables

**Table S1:** Comparison of `scDiagnostics` to existing cell type annotation, OOD detection, and disease-state identification tools.

| Tool Category & Examples | Primary Function | Algorithm-Agnostic? | OOD / Anomaly Detection? | Biological Downstream Analysis? |
|---|---|---|---|---|
| **Standard Reference Mapping** `SingleR` [1], `Azimuth` [2], `CellTypist` [3] | Automated transfer of labels from reference to query datasets. | **No** (Tied directly to their own internal classification algorithms) | **Limited** (Uses relative probability/delta scores; can mask structural shifts) | **No** (Requires manual downstream differential expression) |
| **OOD Detection** `scArches` [5], `eDOC` [38], `scDOT` [39], `scEMAIL` [40] | Latent space mapping designed to reject non-conforming or unknown cells. | **No** (Tied specifically to their internal VAE, Transformer, or Optimal Transport architectures) | **Yes** (Predictive uncertainty, evidential learning, or latent space distance) | **Mixed** (`eDOC` provides gene drivers; most others require manual decoding) |
| **Disease/Perturbation Discovery** `Augur` [30], `MMIL` [41] | Identifies which specific cell types are most responsive to a disease/perturbation. | **N/A** (Does not evaluate annotations; relies on pre-existing labels) | **No** (Assumes cells belong to known clusters and ranks their overall responsiveness) | **Yes** (Ranks cell types by perturbation signature) |
| **Post-Hoc Diagnostics** `scDiagnostics`, `scAnnoX` [42] | Systematic assessment of annotation reliability and structural alignment. | **Yes** (Evaluates labels from any manual or automated workflow) | **Yes** (Absolute structural divergence via Isolation Forests & PCA Reconstruction) | **Yes** (Automated identification of gene shifts driving the anomaly) |

**Table S2:** Annotation confidence scores in $CD14^+$ monocytes classified as anomalous or non-anomalous by `scDiagnostics`. Wilcoxon rank-sum test comparing median prediction/confidence scores across four annotation methods.

| Method | $N_{Anom}$ | $N_{Non\text{-}Anom}$ | Median $Score_{Anom}$ | Median $Score_{Non\text{-}Anom}$ | $p$-value |
|---|---|---|---|---|---|
| `Azimuth` [2] | 3,715 | 3,576 | 0.68 | 0.84 | <0.001 |
| `SingleR` [1] | 3,440 | 3,402 | 0.53 | 0.53 | <0.001 |
| `CellTypist` [3] | 3,851 | 3,594 | 1.00 | 1.00 | <0.001 |
| `scArches` [5] | 3,958 | 3,685 | 0.24 | 0.20 | <0.001 |

**Table S3:** Annotation tool confidence scores in CD14$^+$ monocytes classified as anomalous or non-anomalous by `scDiagnostics` (cell counts listed in Table S2). Wilcoxon rank-sum test comparing median annotation confidence scores (or uncertainty scores for `scArches` [5]) between anomalous and non-anomalous cells across four annotation methods.

| Method | Median Score$_{\text{Anom}}$ | Median Score$_{\text{Non-Anom}}$ | $p$-value |
|---|---|---|---|
| `Azimuth` [2] | 0.68 | 0.84 | <0.001 |
| `SingleR` [1] | 0.53 | 0.53 | <0.001 |
| `CellTypist` [3] | 1.00 | 1.00 | <0.001 |
| `scArches` [5] | 0.24 | 0.20 | <0.001 |

**Table S4:** Concordance between annotation tool confidence-based stratification and `scDiagnostics` anomaly detection in CD14$^+$ monocytes. Spearman rank correlation of $\log_2$-fold changes for 25 interferon signature genes.

| Method | Spearman $r$ | $p$-value |
|---|---|---|
| `Azimuth` [2] | −0.910 | <0.001 |
| `SingleR` [1] | 0.182 | 0.3834 |
| `CellTypist` [3] | −0.642 | <0.001 |
| `scArches` [5] | −0.920 | <0.001 |

**Table S5:** *Col1a2* expression in fibroblasts classified as anomalous or non-anomalous across four annotation methods. Student's $t$-test comparing mean log-normalized *Col1a2* expression between anomalous and non-anomalous cells in DSS-induced colitis samples.

| Method | N$_{\text{Anom}}$ | N$_{\text{Non-Anom}}$ | Mean$_{\text{Anom}}$ | SD$_{\text{Anom}}$ | Mean$_{\text{Non-Anom}}$ | SD$_{\text{Non-Anom}}$ | $p$-value |
|---|---|---|---|---|---|---|---|
| `Azimuth` [2] | 1,060 | 3,337 | 0.212 | 0.672 | 2.109 | 1.196 | <0.001 |
| `SingleR` [1] | 976 | 3,557 | 0.233 | 0.664 | 1.977 | 1.184 | <0.001 |
| `CellTypist` [3] | 2,113 | 3,159 | 0.161 | 0.492 | 1.993 | 1.175 | <0.001 |
| `scArches` [5] | 1,226 | 3,298 | 0.197 | 0.642 | 2.047 | 1.197 | <0.001 |

### References

[1] D Aran, A P Looney, L Liu, E Wu, V Fong, A Hsu, S Chak, R P Naikawadi, P J Wolters, A R Abate, et al. Reference-based analysis of lung single-cell sequencing reveals a transitional profibrotic macrophage. *Nat Immunol*, 20(2):163–72, 2019.

[2] A Butler, C Darby, Y Hao, A Hartman, P Hoffman, G Molla, and R Satija. *Azimuth: a Shiny app demonstrating a query-reference mapping algorithm for single-cell data*, 2023. R package version 0.5.0.

[3] C C Domínguez, C Xu, L B Jarvis, D B Rainbow, S B Wells, T Gomes, S K Howlett, O Suchanek, K Polanski, H W King, et al. Cross-tissue immune cell analysis reveals tissue-specific features in humans. *Science*, 376(6594):eabl5197, 2022.

[4] R Lopez, J Regier, M B Cole, M I Jordan, and N Yosef. Deep generative modeling for single-cell transcriptomics. *Nat Methods*, 15(12):1053–58, 2018.

[5] M Lotfollahi, M Naghipourfar, M D Luecken, M Khajavi, M Büttner, M Wagenstetter, Ž Avsec, A Gayoso, N Yosef, M Interlandi, et al. Mapping single-cell data to reference atlases by transfer learning. *Nat Biotechnol*, 40(1):121–30, 2022.

[6] Z A Clarke, T S Andrews, J Atif, D Pouyabahar, B T Innes, S A MacParland, and G D Bader. Tutorial: guidelines for annotating single-cell transcriptomic maps using automated and manual methods. *Nat Protoc*, 16(6):2749–64, 2021.

[7] T Stuart, A Butler, P Hoffman, C Hafemeister, E Papalexi, W M Mauck III, Y Hao, M Stoeckius, P Smibert, and R Satija. Comprehensive integration of single-cell data. *Cell*, 177(7):1888–902.e21, 2019.

[8] Y Hao, S Hao, E Andersen-Nissen, W M Mauck, S Zheng, A Butler, M J Lee, A J Wilk, C Darby, M Zager, et al. Integrated analysis of multimodal single-cell data. *Cell*, 184(13):3573–87.e29, 2021.

[9] D W Hosmer Jr, S Lemeshow, and R X Sturdivant. *Applied Logistic Regression*, 3rd edition, 2013.

[10] D P Kingma and M Welling. Auto-encoding variational Bayes. In *Proceedings of the 2nd International Conference on Learning Representations*, 2014.

[11] S J Pan and Q Yang. A survey on transfer learning. *IEEE Transactions on knowledge and data engineering*, 22(10):1345–59, 2009.

[12] M Ringnér. What is principal component analysis? *Nat Biotechnol*, 26(3):303–4, 2008.

[13] A J Izenman. *Modern multivariate statistical techniques*. Springer, 2008.

[14] M Woollard, P Panwar, L T G Harland, J Mo, H Zhang, A Swarbrick, B Gottgens, S Ghazanfar, and L H Nghiem. Weighted sliced inverse regression for scalable supervised dimensionality reduction of spatial transcriptomics data. *bioRxiv*, 2025.

[15] G Peyré and M Cuturi. Computational optimal transport: with applications to data science. *Found Trends Mach Learn*, 11(5-6):355–607, 2019.

[16] C Bunne, G Schiebinger, A Krause, A Regev, and M Cuturi. Optimal transport for single-cell and spatial omics. *Nat Rev Methods Primers*, 4:58, 2024.

[17] L Baringhaus and C Franz. On a new multivariate two-sample test. *J Multivar Anal*, 88(1):190–206, 2004.

[18] L Baringhaus and C Franz. Rigid motion invariant two-sample tests. *Stat Sin*, pages 1333–61, 2010.

[19] K V Mardia, J T Kent, and C C Taylor. *Multivariate analysis*. John Wiley & Sons, 2024.

[20] F T Liu, K M Ting, and Z H Zhou. Isolation forest. *Proceedings of the Eighth IEEE International Conference on Data Mining*, pages 413–22, 2008.

[21] L Breiman. Random forests. *Mach Learn*, 45:5–32, 2001.

[22] R A Amezquita, A T L Lun, E Becht, V J Carey, L N Carpp, L Geistlinger, F Marini, K Rue-Albrecht, D Risso, C Soneson, et al. Orchestrating single-cell analysis with Bioconductor. *Nat Methods*, 17(2):137–45, 2020.

[23] A Zeisel, A B Muñoz-Manchado, S Codeluppi, P Lönnerberg, G La Manno, A Juréus, S Marqués, F Mignardi, K Muscato, P Karadzhova, et al. Brain structure. cell types in the mouse cortex and hippocampus revealed by single-cell rna-seq. *Science*, 347(6226):1138–1142, 2015.

[24] E Stephenson, G Reynolds, R A Botting, F J Calero-Nieto, M D Morgan, Z K Tuong, K Bach, W Sungnak, D J Kelvin, M Haniffa, et al. Single-cell multi-omics analysis of the immune response in COVID-19. *Nat Med*, 27:904–16, 2021.

[25] P Cadinu, K N Sivanathan, A Misra, R J Xu, D Mangani, E Yang, J M Rone, K Tooley, Y-C Kye, L Bod, et al. Charting the cellular biogeography in colitis reveals fibroblast trajectories and coordinated spatial remodeling. *Cell*, 187(8):2010–28, 2024.

[26] J Hester and D Vaughan. *bench: High Precision Timing of R Expressions*, 2026. R package version 1.1.4.

[27] D Righelli, L M Weber, H L Crowell, B Pardo, L Collado-Torres, S Ghazanfar, A T L Lun, S C Hicks, and D Risso. SpatialExperiment: infrastructure for spatially-resolved transcriptomics data in R using Bioconductor. *Bioinformatics*, 38(11):3128–31, 2022.

[28] L. Moses and L. Pachter. Museum of spatial transcriptomics. *Nature Methods*, 19:534–546, 2022.

[29] Hervé Pagès, Peter Hickey, and Aaron Lun. *DelayedArray: A unified framework for working transparently with on-disk and in-memory array-like datasets*, 2024. R package version 0.30.1.

[30] M A Skinnider, J W Squair, C Kathe, M A Anderson, M Gautier, K J E Matson, M Milano, T H Hutson, Q Barraud, A A Phillips, et al. Cell type prioritization in single-cell data. *Nature Biotechnology*, 39(1):30–34, 2021.

[31] M D Luecken, S Gigante, D B Burkhardt, R Cannoodt, D C Strobl, N S Markov, L Zappia, G Palla, W Lewis, D Dimitrov, et al. Defining and benchmarking open problems in single-cell analysis. *Nat Biotechnol*, 43(7):1035–40, 2025.

[32] J E Rood, S Wynne, L Robson, A Hupalowska, J Randell, S A Teichmann, and A Regev. The Human Cell Atlas from a cell census to a unified foundation model. *Nature*, 637(8048):1065–71, 2025.

[33] G Heimberg, T Kuo, D J DePianto, O Salem, T Heigl, N Diamant, G Scalia, T Biancalani, S J Turley, J R Rock, et al. A cell atlas foundation model for scalable search of similar human cells. *Nature*, 638(8052):1085–94, 2025.

[34] H Cui, C Wang, H Maan, K Pang, F Luo, N Duan, and B Wang. scGPT: toward building a foundation model for single-cell multi-omics using generative AI. *Nat Methods*, 21(8):1470–80, 2024.

[35] K Z Kedzierska, L Crawford, A P Amini, and A X Lu. Zero-shot evaluation reveals limitations of single-cell foundation models. *Genome Biol*, 26(1):101, 2025.

[36] L Sikkema, M Lkandushi, D Scarcella, A A Moinfar, J Engelmann, and F J Theis. Automated evaluation of single-cell reference atlas mappings enables the identification of disease-associated cell states. *bioRxiv*, 2025.

[37] M Yoshida, K B Worlock, N Huang, R G H Lindeboom, C R Butler, N Kumasaka, C Dominguez Conde, L Mamanova, L Bolt, L Richardson, et al. Local and systemic responses to SARS-CoV-2 infection in children and adults. *Nature*, 602:321–7, 2022.

[38] C Wu, M Zuo, and L Xie. edoc: explainable decoding out-of-domain cell types with evidential learning. *arXiv preprint arXiv:2411.00054*, 2024.

[39] Y-X Xiong and X-F Zhang. scdot: enhancing single-cell rna-seq data annotation and uncovering novel cell types through multi-reference integration. *Briefings in Bioinformatics*, 25(2):bbae072, 2024.

[40] H Wan, L Chen, and M Deng. Scemail: universal and source-free annotation method for scrna-seq data with novel cell-type perception. *Genomics, Proteomics & Bioinformatics*, 20(5):939–958, 2022.

[41] E Craig, T J Keyes, J Sarno, J P D'Silva, P Domizi, M Zaslavsky, A G Tsai, D R Glass, G P Nolan, T Hastie, et al. Annotation-free discovery of disease-relevant cells in single-cell datasets. *Science Advances*, 11(35):eadv5019, 2025.

[42] X Huang, R Liu, S Yang, X Chen, and H Li. scannox: an r package integrating multiple public tools for single-cell annotation. *PeerJ*, 12:e17184, 2024.
